# Clustering-based positive feedback between a kinase and its substrate enables effective T-cell receptor signaling

**DOI:** 10.1101/2020.10.06.328708

**Authors:** Elliot Dine, Ellen H. Reed, Jared E. Toettcher

**Author notes:** Corresponding Author and Lead Contact: Jared E. Toettcher, Lewis Thomas Laboratory Room 140, Washington Road, Princeton, NJ 08544.

## Abstract

Protein clusters and condensates are pervasive in mammalian signaling. Yet how the signaling capacity of higher-order assemblies differs from simpler forms of molecular organization is still poorly understood. Here, we present an optogenetic approach to switch between light-induced clusters and simple protein heterodimers with a single point mutation. We apply this system to study how clustering affects signaling from the kinase Zap70 and its substrate LAT, proteins that normally form membrane-localized clusters during T cell activation. We find that light-induced clusters of LAT and Zap70 trigger potent activation of downstream signaling pathways even in non-T cells, whereas one-to-one dimers do not. We provide evidence that clusters harbor a local positive feedback loop between three components: Zap70, LAT, and Src-family kinases that bind to phosphorylated LAT and further activate Zap70. Overall, our study provides evidence for a specific role of protein condensates in cell signaling, and identifies a simple biochemical circuit that can robustly sense protein oligomerization state.

**Highlights:**

- A general system for studying the role of protein clusters versus dimers.
- Membrane clusters of the kinase Zap70 and its substrate LAT trigger potent downstream signaling.
- Clustering Zap70 with LAT is required for full activation of Zap70 kinase activity.
- A positive feedback loop connects phosphorylated LAT to Zap70 activation via Src-family kinases.

## Introduction

Many cell signaling processes involve the dynamic assembly and disassembly of protein clusters. In some cases, such as Notch/Delta complexes^1^ and death receptor signaling^2^, clusters may emerge due to higher-order oligomerization of the receptor itself upon ligand binding. In others (e.g. receptor tyrosine kinases; the Wnt signalosome), clustering emerges from the convergence of adaptor proteins that bind via modular, multivalent interaction domains to form liquid or gel-like condensates in response to ligand stimulation^3^. Recent advances in imaging have established that that protein clustering can accompany signaling pathway activation *in vivo*^4–6^, and biochemical reconstitution experiments demonstrate kinase-triggered clustering of minimal sets of components *in vitro*^1–9^, suggesting that mesoscale protein assemblies are fundamental to eukaryotic cell signaling.

Yet a key question remains: do clusters play an active role in shaping signaling responses, or do they simply emerge as an unavoidable byproduct of the weak multivalent interactions that occur between signaling proteins^10^? Discriminating between these two possibilities presents a challenging problem. The protein-protein interaction domains that typically drive clustering also perform other essential signaling functions (e.g., recruiting enzymes to their substrates); therefore, one cannot just delete these domains and assume that the resulting signaling deficiencies are caused by a loss of clustering. The recent development of chemical biology and optogenetic tools for inducing protein clustering offers a potential solution^11–17^. By triggering the assembly of clusters containing selected proteins of interest and comparing to other forms of molecular interaction (e.g., 1-to-1 heterodimers), we might directly test for the functional consequences of clustering. Such user-defined “signalosomes” could also prove useful to the synthetic biologist, enabling the clustering of defined proteins to confer specific signal processing functions^18,19^.

Here, we define the biochemical function of protein clustering in one specific cellular context: the phase separation of two proteins, Zap70 and Linker of Activated T-cells (LAT), that are essential for linking the activation of the T cell receptor to downstream signaling pathways. Following activation of the T cell receptor, the kinase Lck phosphorylates Zap70, which goes on to phosphorylate the membrane-tethered scaffold protein LAT^20^. LAT is a tyrosine rich protein that, when phosphorylated, can undergo liquid-liquid phase separation due to interactions with other multivalent signaling proteins: Grb2, SOS, and PLCγ^7,9,21^ (**Figure 1A)**. It was previously shown that three tyrosine-to-phenylalanine mutations on LAT is sufficient to abolish both clustering and prevent Erk activation and intracellular calcium release. However, LAT tyrosines are also needed to recruit Grb2 and PLCγ to the membrane, without which downstream signaling cannot proceed^22^, limiting our ability to understand the functional role of LAT clustering from this type of experiment.

**Figure 1:**
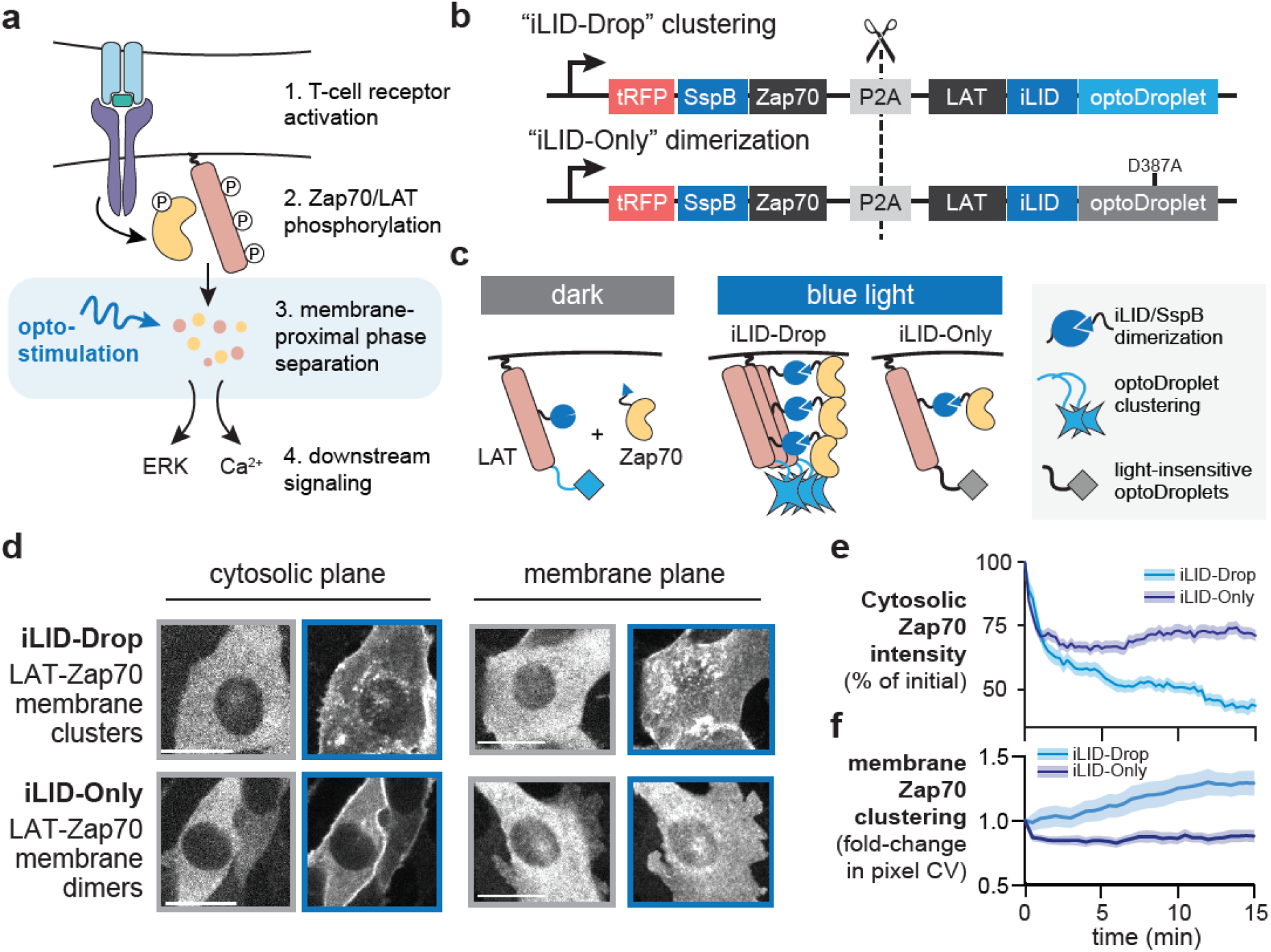
Development of optogenetic systems to compare Zap70:LAT oligomerization states. (**a**) Cartoon depicting the known TCR pathway and the role of protein phosphorylation and Zap70:LAT clustering in activating downstream signaling pathways. In this study, we use optogenetic tools to plug in at the step of Zap70:LAT clustering and see how different forms of the Zap70:LAT interaction affect downstream signaling. (**b**) Design of the optogenetic constructs to compare dimerization and clustering of Zap70 and LAT. iLID-Only construct contains a mutation in Cry2 (D387A) that prevents light-dependent homo-oligomerization; tRFP=TagRFP. (**c**) Cartoon of what occurs upon light stimulation of the optogenetic constructs shown in **b**. Adapted from Ref. 20. (**d**) Images of Zap70 tagged with TagRFP localization in NIH-3T3 cells taken at two different planes with spinning disk confocal imaging. Gray border indicates images taken prior to blue light illumination, and blue border indicates images taken following 5 minutes of stimulation. Scale bars = 20 μm. Note: because TagRFP brightness is increased by blue light illumination, images were auto-scaled independently before and after light stimulation. (**e**) Quantification of the change in cytosolic fluorescence across 15 minutes of blue light illumination in both iLID-Only and iLID-Drop cells. n ≥ 20 cells in both conditions. (**f**) Quantification in change of the coefficient of variation (CV) of TagRFP intensity for images taken in the membrane plane during 15 minutes of blue light illumination. n = 20 cells in both conditions.

Our aim was to precisely define the contribution of protein clustering in a minimal T cell activation module – the interaction between the kinase Zap70 and its substrate LAT – to relate the formation of specific molecular assemblies to the cell’s resulting signaling state (**Figure 1A)**. We engineered optogenetic variants of Zap70 and LAT whose 1-to-1 dimerization or assembly into droplets could be switched with a single point mutation, and tested their sufficiency for signaling in fibroblasts that lack other T cell-specific signaling components to eliminate other clustering stimuli (e.g., clusters of the T cell receptor itself). Remarkably, Zap70:LAT clusters were fully competent to trigger downstream Erk and calcium signaling, whereas Zap70:LAT heterodimers produced no signaling response. Subsequent experiments and computational modeling revealed that clustering-induced signaling requires a 3-component positive feedback loop between Zap70, its substrate LAT, and a Src-family kinase whose recruitment to LAT enables further activation of Zap70. Our results suggest that the dual ability of Src to bind phospho-tyrosines and phosphorylate nearby proteins forms a circuit that is highly sensitive to changes in protein oligomerization state, providing a robust clustering-based signaling switch that may be broadly used by endogenous signaling systems and could also find application in synthetic kinase-based circuits.

## Results

### An optogenetic platform for directing Zap70:LAT dimerization versus condensation

Our first goal was to create optogenetic tools that could be used to acutely trigger distinct modes of interaction between Zap70 and LAT: forming either one-to-one Zap70:LAT heterodimers or higher-order clusters of heterodimers upon illumination. Ideally, such a system would enable the experimentalist to toggle between dimers or cluster-of-dimers without changing any other parameters of the system (**Figure 1B-C**). To accomplish this goal, we outfitted LAT with two optogenetic systems to independently control its dimerization with Zap70 versus assembly into higher-order clusters. For Zap70:LAT dimerization, we turned to the iLID-SspB system^23^, which forms one-to-one heterodimers with a binding affinity of ~100 nM in response to blue light^23,24^. For LAT clustering, we took advantage of the optoDroplet system, which can be used to trigger membrane-localized protein droplets upon blue light stimulation^16,17^. Crucially, a single point mutation in the Cry2 component of optoDroplets (Cry2 D387A^25^) renders it completely insensitive to blue light, preventing cluster formation^13^. We thus engineered two DNA constructs: one that expresses that expresses a LAT-iLID-optoDroplet fusion protein with a fluorescent, SspB-tagged Zap70 (termed “iLID-Drop”), and one that is identical except for the light-insensitive point mutation in the optoDroplet system (termed “iLID-Only”) (**Figure 1B,C**). We reasoned that this matched pair of systems constituted an ideal test case because Zap70:LAT dimerization would be controlled by identical iLID-SspB interactions both cases, with only the additional clustering of Zap70:LAT heterodimers depending on the functionality of the optoDroplet system.

We initially set out to test whether the iLID-Drop and iLID-Only tags could indeed drive different forms of Zap70:LAT interactions. We transduced NIH-3T3 cells with lentiviral vectors expressing one or the other, sorted them for the same TagRFP levels to ensure closely-matched expression in both cell lines (**Figure S1A-C**), and imaged the resulting cell lines by confocal microscopy. We observed rapid cytosolic depletion of TagRFP fluorescence upon illumination in both iLID-only and iLID-Drop cells, consistent with recruitment of cytosolic TagRFP-SspB-Zap70 to membrane-localized LAT-iLID (**Figure 1D-E, Movie S1**). Only iLID-Drop cells exhibited nucleation and growth of small membrane-localized TagRFP clusters (**Figure 1D; Movie S1-2**), an effect that could be quantified by measuring the variance of TagRFP pixel intensities at the plasma membrane over time (**Figure 1F**). Despite similar initial kinetics of cytosolic depletion between cell lines, we observed some additional cytosolic depletion of Zap70 in iLID-Drop cells on the same timescale as membrane clustering (**Figure 1E**), suggesting that Zap70:LAT clusters may increase Zap70’s retention at the membrane as previously observed on supported lipid bilayers^26,27^. Nevertheless, any differences in Zap70 cytosolic depletion were minor compared to the variability in expression levels between cells (compare cells in **Figure 1D**; lower-left panels). Overall, our results indicate that both the iLID-Drop and iLID-only systems recruit Zap70 to LAT, but only iLID-Drop induces the formation of membrane-localized Zap70:LAT clusters. These differences in molecular organization are achieved using a single point mutation and at identical expression levels, thereby providing a controlled platform for assessing the functional consequences of clustering.

### Zap70:LAT clusters but not heterodimers activate downstream signaling pathways

How does dimerization vs clustering of LAT and Zap70 affect the activation of downstream signaling pathways? To address this question, we set out to monitor downstream signaling in iLID-Only and iLID-Drop fibroblasts. Fibroblasts are an ideal cellular context for this study, as they lack T cell-specific components that could trigger clustering independently of our optogenetic systems^28^, but still harbor intact downstream MAPK and calcium signaling for monitoring downstream cellular responses. We thus expressed the iLID-Drop and iLID-Only systems in NIH-3T3 mouse fibroblasts that were also engineered to express live-cell biosensors of downstream signaling: the Erk Kinase Translocation Reporter (ErkKTR) and GCaMP6f **(Figure 2A-B)**. The ErkKTR leaves the nucleus upon activation of Erk signaling^29^, while GCaMP6f (GCaMP) becomes much brighter upon release of Ca^2+^ from stored vesicles^30^.

**Figure 2:**
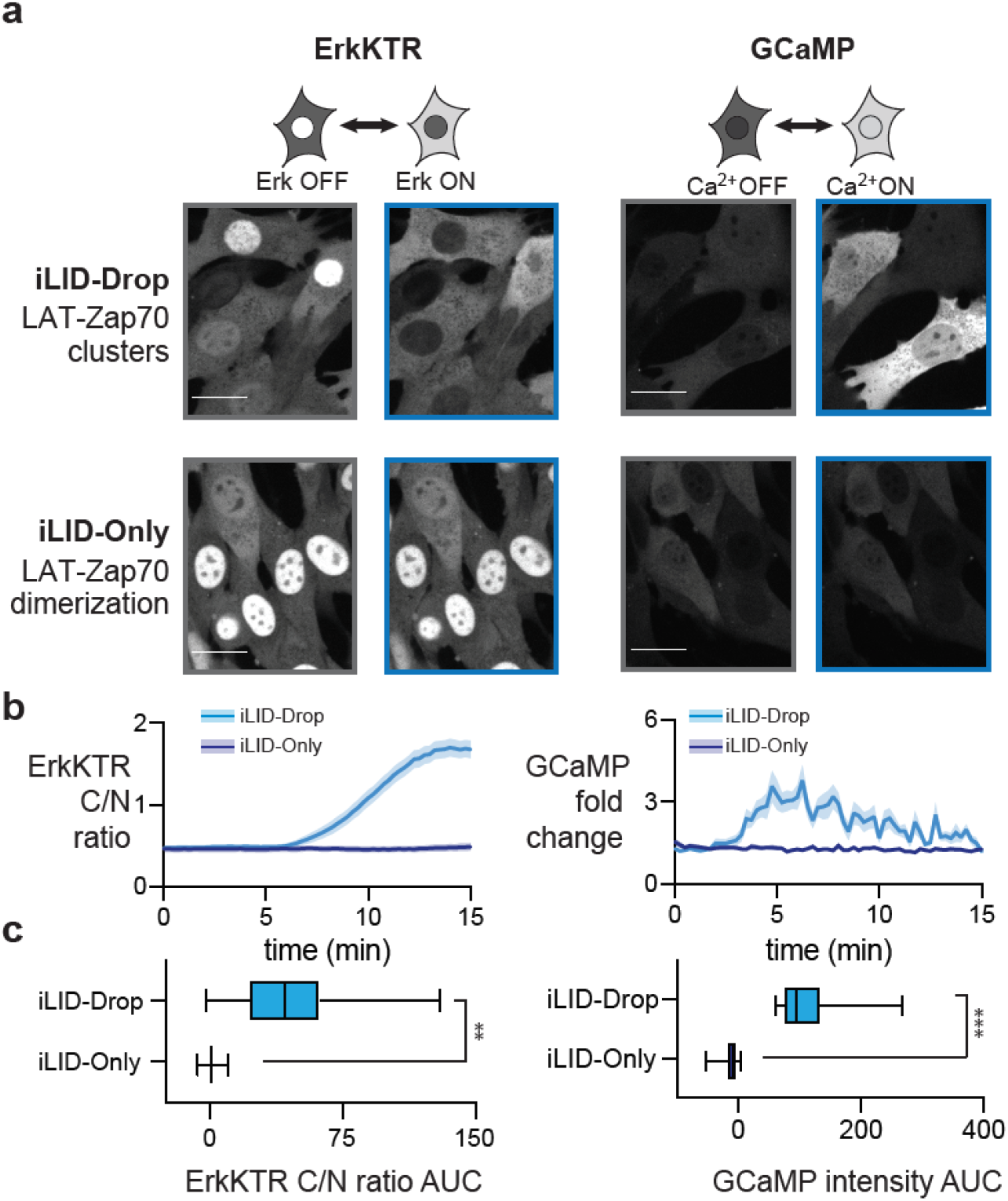
Clustering but not heterodimerization of Zap70 and LAT induces signaling. (**a**) Images of ErkKTR-irFP and GCaMP in NIH3T3 cells expressing iLID-Only and iLID-Drop constructs. Images are representative of pre- and post-stimulation responses. Scale Bar = 20 μm. (**b-c**) Quantification of cytosolic to nuclear (C/N) ratio for ErkKTR irFP and GCaMP fluorescence fold change for iLID-Only and iLID-Drop expressing cells. Data shows time courses (in **b**) and area under the curve (in **c**) for n ≥ 50 cells from 4 different experiments for each cell line. For **c**, boxes represent 25th - 75th percentile and whiskers show the minimum and maximum values. Statistical significance computed from 4 independent experiments using Student’s T test with ** = p < 0.01 and *** = p < 0.001.

We first generated a single NIH-3T3 cell line expressing both irFP-tagged ErkKTR and GCaMP, and then transduced and sorted for identical expression levels of either our red fluorescent Zap70:LAT ILID-Drop and ILID-Only constructs (**Figure S1C**). Both cell lines were plated, washed and starved in serum-free media for 2 hours, and monitored for Erk and calcium responses after blue light stimulation (**Figure 2A**). Light-stimulated iLID-Drop cells exhibited near-complete export of ErkKTR-irFP from the nucleus and repeated spikes of GCaMP fluorescence, indicative of strong Erk and calcium signaling responses, within minutes after blue light illumination (**Figure 2A-B, Movie S3**). No such responses were observed in iLID-Only cells, despite similar light-induced translocation of Zap70 to the cell membrane (**Figure 2A-B, Movie S4**). We used the area under the curve (AUC) of biosensor activity in each cell to quantify and compare responses, revealing significant increases for both Erk and calcium signaling in iLID-Drop cells as compared to iLID-Only cells (**Figure 2C**). Taken together, our data indicates that membrane-localized clusters of Zap70 and LAT are sufficient to trigger Erk and calcium signaling responses even in non-T cells, whereas Zap70:LAT heterodimers do not.

### Phosphorylation and activation of Zap70 is the key clustering-dependent step

We next sought to identify the biochemical steps that are activated by Zap70:LAT clustering to trigger downstream signaling. Membrane clusters have been suggested to play many distinct and separable functions, such as enhancing reaction rates by increasing local concentration, excluding negative regulators to locally increase the levels of phosphorylated species, or even altering the processivity of a kinase for its substrate to drive efficient multi-site phosphorylation^31,33^. As a first step towards identifying the mechanism for clustering-induced signaling, we monitored each of the steps normally associated with Zap70/LAT activation (**Figure 3A**). During T cell activation, the Zap70 kinase is first activated by phosphorylation at Tyr319. Activated Zap70 then phosphorylates LAT on four sites, three of which (Tyr171, Tyr191 and Tyr226) are rapidly phosphorylated and one of which (Tyr132) is phosphorylated more slowly and has been proposed to serve as the kinetic proofreading step for responding only to high-affinity TCR-ligand interactions^22,34–36^.

**Figure 3.**
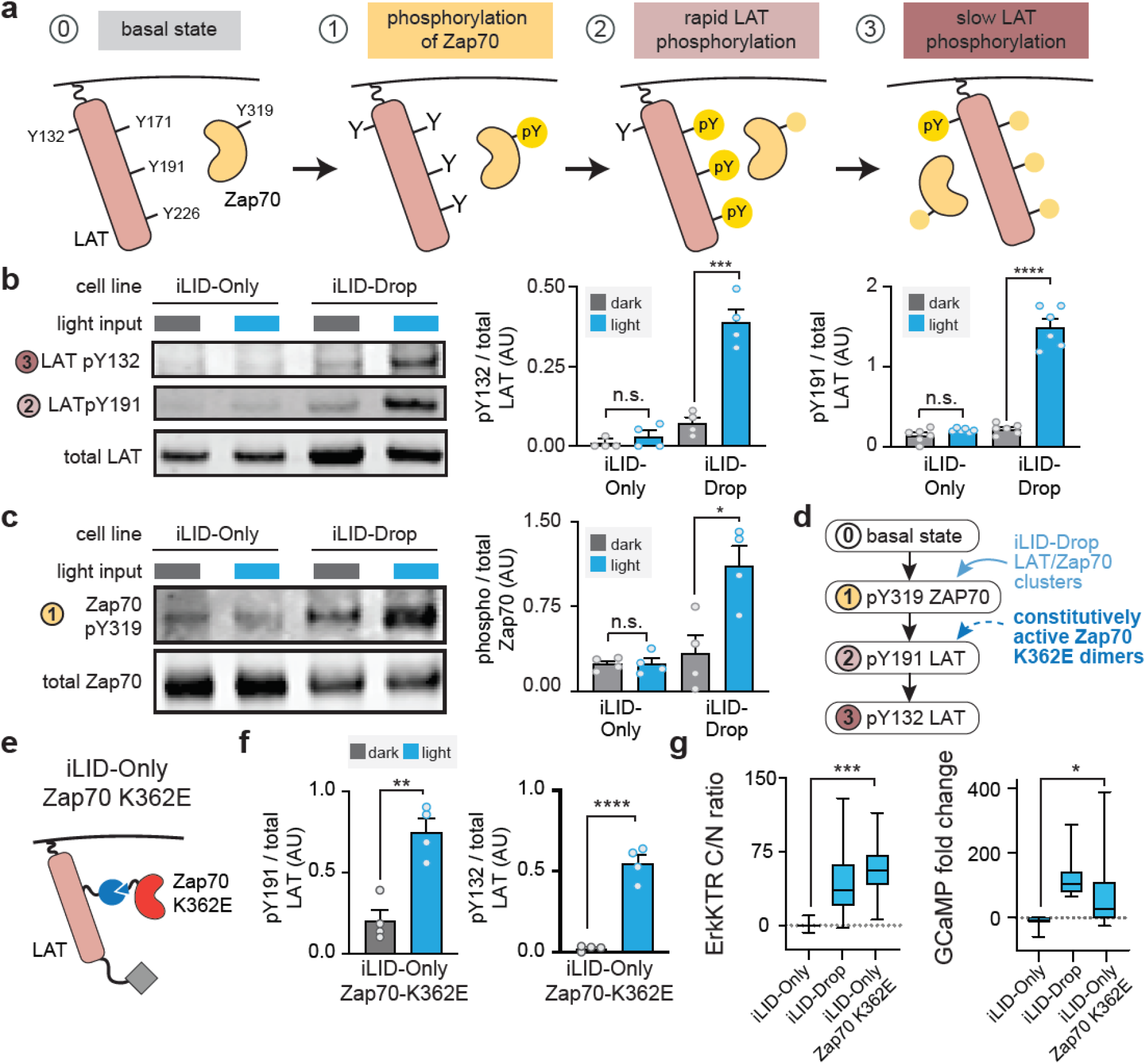
Clustering is required for light-induced Zap70 and LAT phosphorylation. (**a**) Cartoon representing the three steps of Zap70:LAT interaction. Zap70 is first activated by phosphorylation on Tyr319, and then rapidly phosphorylates LAT at Tyr171, 191, and 226. Finally, Zap70 slowly phosphorylates Tyr132 on LAT. (**b**) Western blot and quantification of phospho-LAT at positions Y132 and Y191 in the dark and after 20 minutes of blue light stimulation. (**c**) Western blot and quantification of pY319-Zap70 in the dark and after 20 minutes of blue light stimulation. (**d**) Schematic of the LAT-Zap70 cascade showing expected modes of action of two optogenetic control schemes: clustering of LAT and Zap70 versus dimerization between LAT and Zap70^K362E^ In the latter case, the activating Zap70 mutation would bypass the requirement for Zap70 phosphorylation, potentially enabling LAT phosphorylation from dimers alone. (**e**) Cartoon of iLID-Only Zap70 K362E. (**f**) Quantification of Western blots for pY191 and pY132 LAT in cells expressing iLID-Only Zap70 K362E. (**g**) Quantification of the integrated area-under-the-curve of signaling responses from ErkKTR-irFP (C/N ratio) and GCaMP (fold change). Boxes represent 25th – 75th percentile and whiskers show minimum and maximum values. n ≥ 30 data points are shown from 3 different experiments for all cell lines. Graphs display mean ± SEM and independent biological replicates (points). All statistical comparisons were performed using the Student’s T test using all independent biological replicates. * = p < 0.05, ** = p < 0.01, *** = p < 0.001 and **** = p < 0.0001.

To test which steps in the Zap70:LAT cascade were dependent on light-induced clustering, we quantified Zap70 Tyr319, LAT Tyr191, and LAT Tyr132 phosphorylation under dark and illuminated conditions. We found that all three sites were phosphorylated in a light-dependent manner in iLID-Drop cells but not in iLID-Only cells, suggesting that clustering is required even for the top-most phosphorylation event in the Zap70:LAT cascade: activation of Zap70 itself (**Figure 3B-C**). We further confirmed that clustering-specific phosphorylation of Zap70, LAT, and downstream signaling proteins could be observed in human-derived HEK-293T cells, demonstrating that our results were not specific to a single cell line and applied to cells of both mouse and human origin (**Figure S2)**. Finally, to confirm that Zap70’s Tyr319 phosphorylation is indeed necessary for downstream signaling, we mutated this residue to phenylalanine, and found that it abolished downstream signaling in illuminated iLID-Drop cells (**Figure S3A-B**), consistent with prior reports that phosphorylation at this residue is required for Zap70 activation and downstream signaling^37,38^.

Our data indicates that clustering is required for the initial step of Zap70 phosphorylation and activation, but is this its only role? We reasoned that if clustering is only required for Zap70 activation, then a constitutively-active Zap70 variant should be able to elicit a full signaling response even in iLID-Only cells. We thus established an iLID-Only NIH-3T3 cell line using a previously characterized Zap70 allele, Zap70^K362E^, that exhibits weak constitutive activity even in the absence of its phosphorylation^39^ (**Figure 3E**). Light stimulation of iLID-Only Zap70^K362E^ cells triggered LAT phosphorylation at all tyrosine residues tested (**Figure 3F**), and we observed downstream signaling that generally matched what was observed in iLID-Drop cells (**Figure 3G, Movie S5**). Taken together, these data demonstrate that light-induced clustering is required for initiation of signaling in the minimal Zap70/LAT module, and that the requirement for clustering can be bypassed by providing a constitutively active Zap70. These data also constitute an important control, ruling out the possibility that the iLID-Only system is incapable of triggering downstream signaling – for example, if the complexes induced by iLID-SspB dimerization were somehow incapable of supporting Zap70-to-LAT phospho-transfer.

### Clustering-induced Zap70 activation requires both kinase activity and substrate residues

What occurs within Zap70:LAT clusters to promote Zap70 phosphorylation? To gain insight into this process, we set out to establish the requirements for clustering-based signaling using LAT and Zap70 mutant variants. We first tested whether Zap70 kinase activity is required by constructing an iLID-Drop variant containing a kinase-dead Zap70 mutant -- Zap70 K369R^40^. This kinase-dead variant failed to induce Zap70 phosphorylation, even though illumination still produced membrane-associated Zap70:LAT clusters, indicating that Zap70 phosphorylation depends on Zap70 kinase activity (**Figure 4A-B**). We also tested whether Zap70 phosphorylation required the presence of LAT as a substrate. We thus engineered an iLID-Drop variant in which LAT was replaced by a variant (LAT^FFF^) lacking the tyrosines that serve as the first substrates for Zap70 phosphorylation^7,9,22^. Again, no light-induced increase in Zap70 phosphorylation was observed in LAT^FFF^ iLID-Drop cells, despite light-induced Zap70 membrane localization and clustering (**Figure 4C-D**). iLID-Drop variants harboring each single Y-to-F mutation in LAT still robustly triggered downstream signaling (**Figure S3C**), as has been observed in T cells, suggesting that the requirement is not restricted to any single Tyr residue^22^.

**Figure 4.**
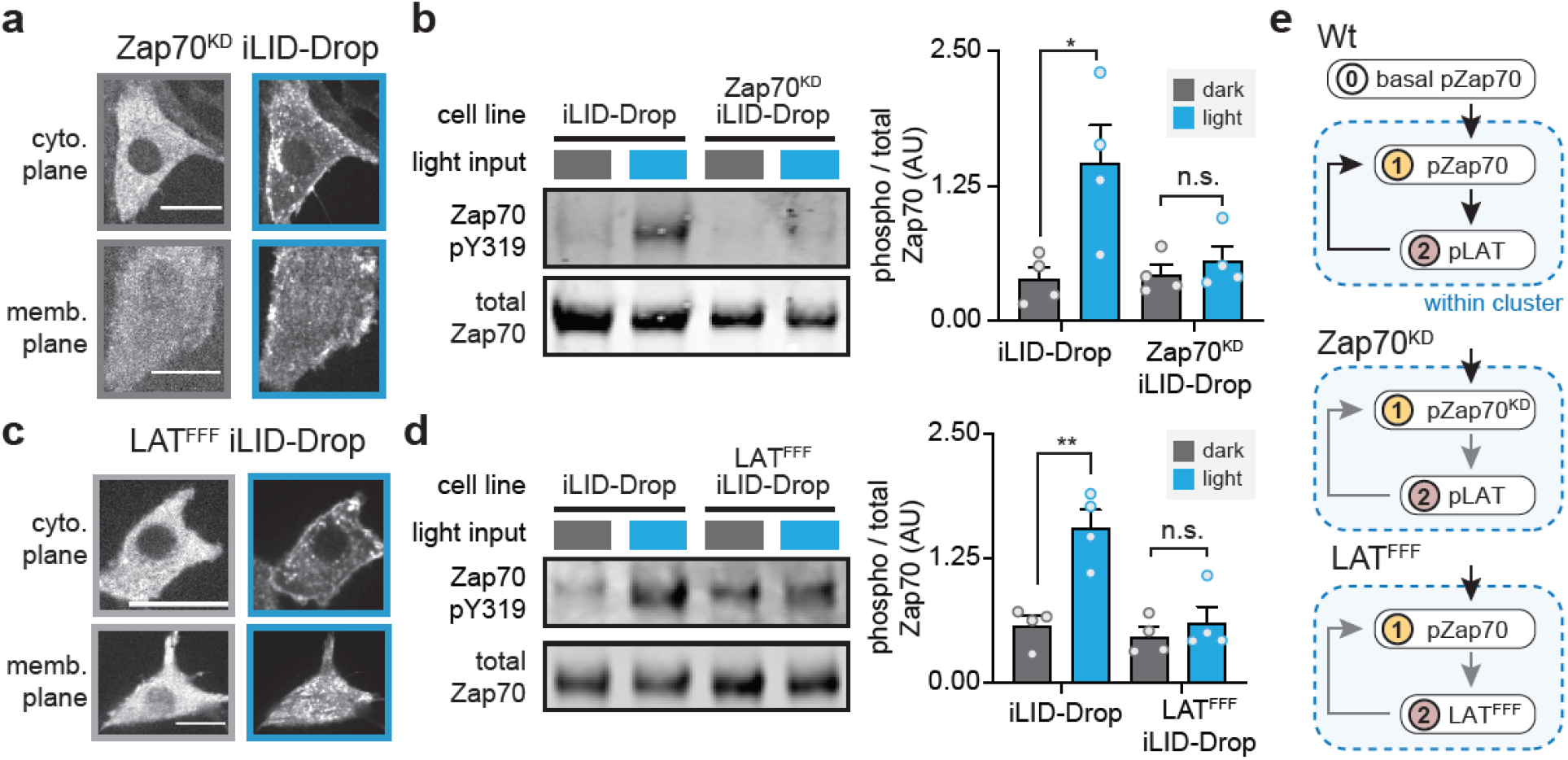
Positive feedback links Zap70 kinase activity and LAT substrate phosphorylation with further Zap70 activation. (**a**) Images of TagRFP-labeled, kinase-dead Zap70^K369R^ mutant (Zap70^KD^) in iLID-Drop cells. Images show cytosolic and membrane planes to capture both clustering and cytosolic depletion of Zap70, before and after light stimulation (blue and gray borders, respectively). Scale bars = 20 μm. (**b**) Western blot and quantification of pY319-Zap70 in the dark and after 20 minute of blue light stimulation for wild-type iLID-Drop and Zap70KD iLID-Drop. (**c**) Images of TagRFP-Zap70 localization in LAT^FFF^ (LAT Y171F, Y191F, Y226F) iLID-Drop cells. Images were taken as in **a**. (**d**) Western blot and quantification of pY319-Zap70 in the dark and after 20 minutes of blue light stimulation for wild-type and LAT^FFF^ iLID-Drop cells. (**e**) Cartoon of positive feedback loop that occurs between LAT and Zap70 phosphorylation inside the iLID-Drop clusters. No increase in pZap70 is observed in either Zap70^KD^ or LAT^FFF^ cells, demonstrating that this upstream even depends on downstream steps, a diagnostic sign of positive feedback. Note for panels **a** and **c**: because the brightness of TagRFP is increased by blue light illumination, images were auto-scaled separately before and after light stimulation. Graphs display mean ± SEM and independent biological replicates (points). All statistical comparisons were performed using the Student’s T test using all independent biological replicates. * = p < 0.05, ** = p < 0.01, *** = p < 0.001 and **** = p < 0.0001.

Taken together, our data shows that only clusters containing catalytically active Zap70 and phosphorylatable LAT can be fully activated. The dependency of an upstream event (Zap70 phosphorylation) on downstream attributes (Zap70 kinase activity and a phosphorylatable LAT substrate) is indicative of a positive feedback loop operating within Zap70:LAT clusters (**Figure 4E)**. This feedback loop may operate as follows: a low amount of basally-phosphorylated Zap70 phosphorylates LAT within the cluster, which – through an as-yet-undefined mechanism – triggers additional phosphorylation and activation of Zap70. Fully-active Zap70 further phosphorylates LAT, culminating with the activation of downstream signaling pathways.

### Src-family kinases implement feedback linking LAT phosphorylation to Zap70 activation

We next sought to identify the kinase that mediates Zap70 phosphorylation within membrane-associated Zap70:LAT clusters. During T cell activation, Zap70 Tyr319 is phosphorylated by the Src-family kinase Lck^37,41^. Although NIH-3T3 and HEK-293T cells do not normally express Lck, they do possess general-purpose Src-family kinases (Src, Yes and Fyn) that we reasoned could play an analogous role in illuminated iLID-Drop cells. While Src-family kinases are required for establishing an initial pool of phosphorylated, active Zap70, whether they play additional roles in membrane-localized signaling clusters independently of the T cell receptor remains unclear.

We began by testing whether light-induced Zap70 phosphorylation in NIH-3T3 cells depends on Src-family kinase (SFK) activity using the small-molecule kinase inhibitors PP1 or PP2 to inhibit SFK activity (**Figure 5A)**. Indeed, we observed that these inhibitors eliminated Zap70 Tyr319 phosphorylation in all conditions (**Figure 5B**). For cleaner control over SFK activity, we next expressed the iLID-Drop system in “SYF” mouse embryonic fibroblasts that were engineered to lack all three ubiquitous SFKs, Src, Yes and Fyn^42^. Just as in the PP1/PP2 experiments, we found that clustering-induced Zap70 phosphorylation was completely abolished in SYF fibroblasts regardless of illumination conditions but was fully restored by expression of Src (**Figure 5C-D)**. This restoration required Src kinase activity, as SYF iLID-Drop cells expressing a kinase-dead Src allele (Src K297R^42^) also failed to produce Zap70 phosphorylation **(Figure 5D)**.

**Figure 5.**
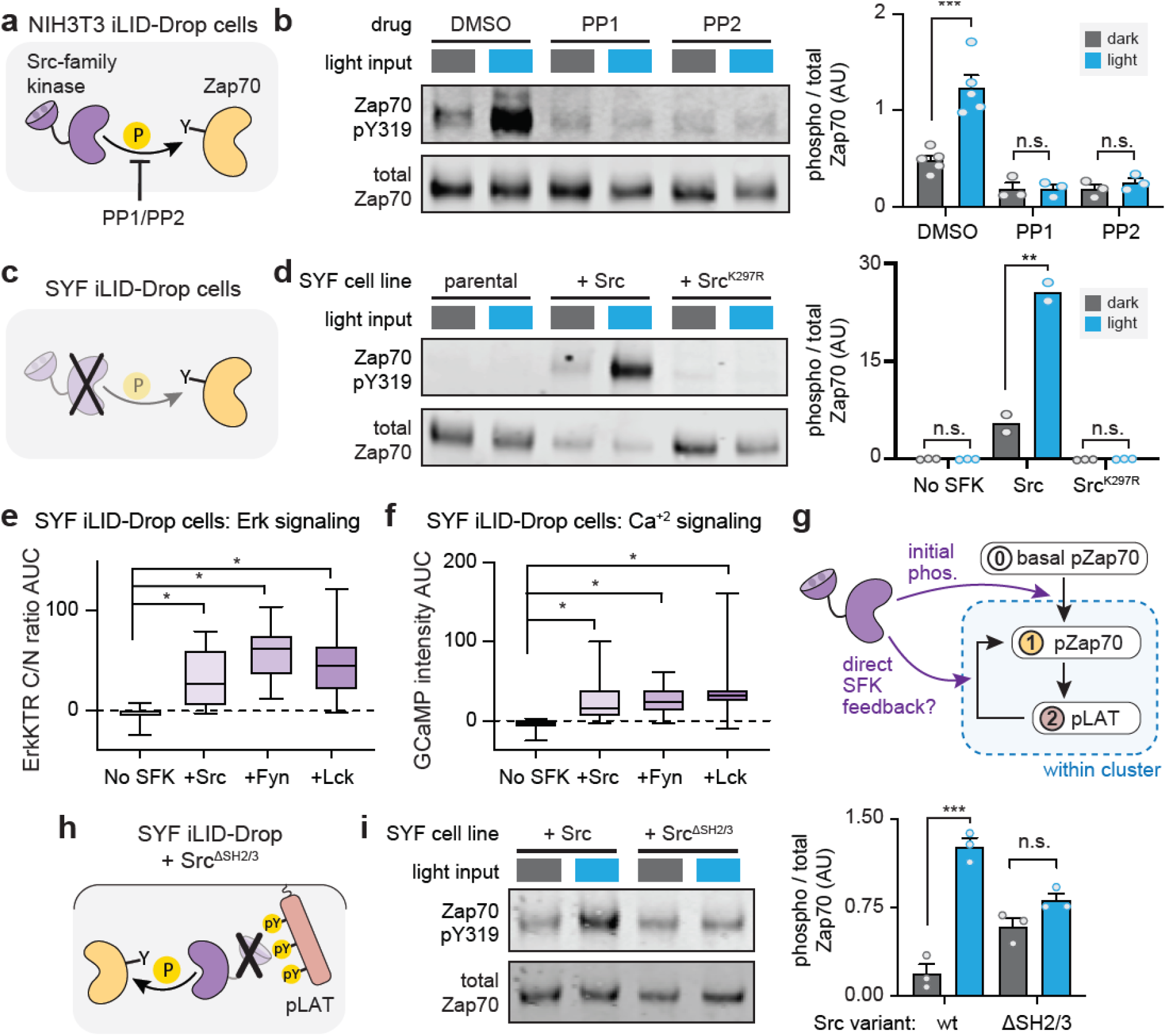
Positive feedback driven Zap70 activation depends on Src-family kinase (SFK) activity. (**a**) Schematic of treatment with the SFK inhibitors PP1 or PP2 in NIH-3T3 iLID-Drop cells. (**b**) Western blot and quantification of phospho-Zap70 after 20-minute treatment with PP1 and PP2 versus DMSO. Results from dark (gray) and light-stimulated (blue) cells are shown. (**c**) Schematic of experiments in iLID-Drop SYF cells. (**d**) Western blot and quantification of pY319-Zap70 in parental SYF cells or SYF cells expressing Src or a kinase-dead Src allele (Src^K297R^). (**e-f**) Area under curve of ErkKTR-irFP (C/N ratio; in **e**) or GCaMP (fold change; in **f**) for iLID-Drop SYF cells. Boxes represent 25th – 75th percentile and whiskers show minimum and maximum values. n ? 20 cells from 2 different experiments. (**g**) Schematic showing two steps in the LAT-Zap70 feedback circuit where SFK activity may be required. (**h**) Schematic of experiments in Src^ΔSH2-3^ iLID-Drop SYF cells. (**i**) Western blot and quantification of pY319-Zap70 in SYF MEFs with Src or Src^ΔSH2-3^ added back. Graphs display mean ± SEM and independent biological replicates (points). All statistical comparisons were performed using the Student’s T test using all independent biological replicates. * = p < 0.05, ** = p < 0.01, *** = p < 0.001 and **** = p < 0.0001.

To further probe the generality of our results, we characterized the dependence of clustering-induced signaling on the identity and expression levels of the Src-family kinases present in our experiments. We first tested whether any of three different SFKs (Src, Fyn or Lck) were similarly capable of rescued clustering-induced signaling. Indeed, we found that iLID-Drop SYF cells expressing Src, Fyn or Lck triggered similar levels of Erk and calcium signaling (**Figure 5E-F**). Second, we compared the expression levels of our Src-transduced SYF cells to endogenous Src expression in NIH-3T3 cells by Western blotting. We observed that Src-transduced SYF cells expressed ~100-fold higher levels of Src than NIH-3T3s cells (**Figure S4A-B**). Nevertheless, pathway activation still robustly depended on cluster formation, as iLID-Only expressing SYF-MEFs failed to mount a signaling response (**Figure S4C**). Taken together, our data demonstrate that Src-family kinase activity is essential for Zap70 activation and LAT phosphorylation in NIH-3T3 cells, and that this effect appears to be robust across different Src-family kinase family members and a wide range of their expression levels.

Based on our data and classic studies of Zap70 activation^37,43^, we envisioned two potential roles that Src-family kinases might play in our system. First, leaky SFK activity may be required to provide an initial basal level of Zap70 phosphorylation, an effect that we observed in dark iLID-Drop and iLID-Only cells throughout our study (**Figure 3C** an **5D**). This leaky activity may be a prerequisite for initial Zap70 phosphorylation of LAT, which might then be amplified by SFK-independent positive feedback to generate full phosphorylation of Zap70. Second, SFKs may also *directly* participate in positive feedback by binding to phospho-LAT and then phosphorylating nearby Zap70 molecules, further increasing Zap70 activity and LAT phosphorylation (**Figure 5G)**. This second possibility is supported by structural studies of Src activation: Src contain an SH2 domain that can lock it in an auto-inhibited conformation until it binds to pTyr residues, which both tethers Src to a potential substrate and increases its activity^44^. The binding of an SFK’s SH2 domain to LAT’s pTyrs, possibly strengthened further by binding between the SFK’s SH3 domain and a proline-rich motif on LAT^39^, could thus trigger recruitment and local activation of Src-family kinases within the cluster, driving further Zap70 and LAT phosphorylation in a positive feedback loop^45,46^.

To separate these two potential functions of SFKs, we set out to introduce a “feedback-disconnected” Src variant that could still drive basal Zap70 phosphorylation but not participate in positive feedback within Zap70:LAT clusters. To do so we deleted the SH2 and SH3 domains from our previously made BFP-tagged Src (Src^ΔSH2/3^-BFP). This Src variant should lack all autoinhibitory interactions and so exhibit high activity, supporting basal Zap70 phosphorylation. However, it should also lack any protein association domains for recruitment to phospho-LAT, thereby blocking any potential role in cluster-localized positive feedback (**Figure 5H**). We engineered iLID-Drop SYF fibroblasts to express either Src^ΔSH2/3^-BFP or Src-BFP at levels that resulted in basal Zap70 phosphorylation in the dark, and tested both cell lines for an increase in Zap70 phosphorylation upon light stimulation. As before, we found that Zap70 phosphorylation increased upon light stimulation in Src-BFP iLID-Drop cells (**Figure 5I**). This effect was dramatically reduced in Src^ΔSH2ΔSH3^-BFP iLID-Drop cells, which showed similar levels of phosphorylation in both dark and light and failed to attain the high levels of Zap70 phosphorylation observed in light-stimulated Src-BFP cells (**Figure 5I**). We thus conclude that Src is not simply required to establish basal Zap70 phosphorylation, but is also directly involved in positive feedback upon light stimulation in a manner that depends on its SH2 and SH3 protein association domains.

Our data can be readily interpreted in the context of a simple conceptual model: a cluster-localized positive feedback loop involving Zap70, LAT and an SFK. Basally phosphorylated Zap70 leads to weak LAT phosphorylation, followed by SFK recruitment and activation through SH2-mediated binding to pLAT. The SFK then further phosphorylates nearby Zap70 proteins within the cluster, completing the feedback loop. Strikingly, the system appears to be a high-fidelity sensor of clustering state, with all-or-none signaling differences observed between clustered and un-clustered LAT, even when the identity or expression level of the Src-family kinase is varied.

### A mathematical model recapitulates signaling through cluster-localized positive feedback

The presence of feedback can make it extremely difficult to intuit the behavior of a biochemical network, even when such a system consists of only three components^47^. We thus wondered whether we could recapitulate our experimental observations – including the responses observed from clusters, dimers, and various mutant proteins – using a minimal mathematical model of the three-component signaling circuit. We reasoned that such a model could be tested for its sufficiency to recapitulate our experimental observations and to explore additional scenarios for further insights into the Zap70/LAT/SFK module.

Our mathematical model contains three proteins (LAT, Zap70 and Src) that can occupy two cellular compartments: a cytosolic compartment containing free Zap70 and Src, and a membrane-localized compartment containing LAT, bound Zap70:LAT, and bound Src:p-LAT (**Figure 6A**). Our model incorporates two light-dependent effects. First, we assume that Zap70 has an increased propensity to phosphorylate LAT when the two proteins are tethered by light-induced iLID-SspB dimerization. Second, we model the light-induced formation of LAT clusters as a simple decrease in their available volume, thus leading LAT and any LAT-bound proteins to become proportionally concentrated in the cluster. We model two binding interactions (light-induced Zap70:LAT binding through the iLID-SspB interaction^23^; Src:p-LAT binding through its SH2 domain^48^) and three phosphorylation reactions (weak Zap70 phosphorylation by free Src; strong Zap70 phosphorylation by Src:p-LAT complexes; and LAT phosphorylation by p-Zap70). Finally, we assume simple, first-order dephosphorylation of LAT and Zap70 by ubiquitous phosphatases in the cell. Where possible, we inferred model parameters from experimental measurements of the relevant proteins (**Supplementary Methods, Table S1**).

**Figure 6.**
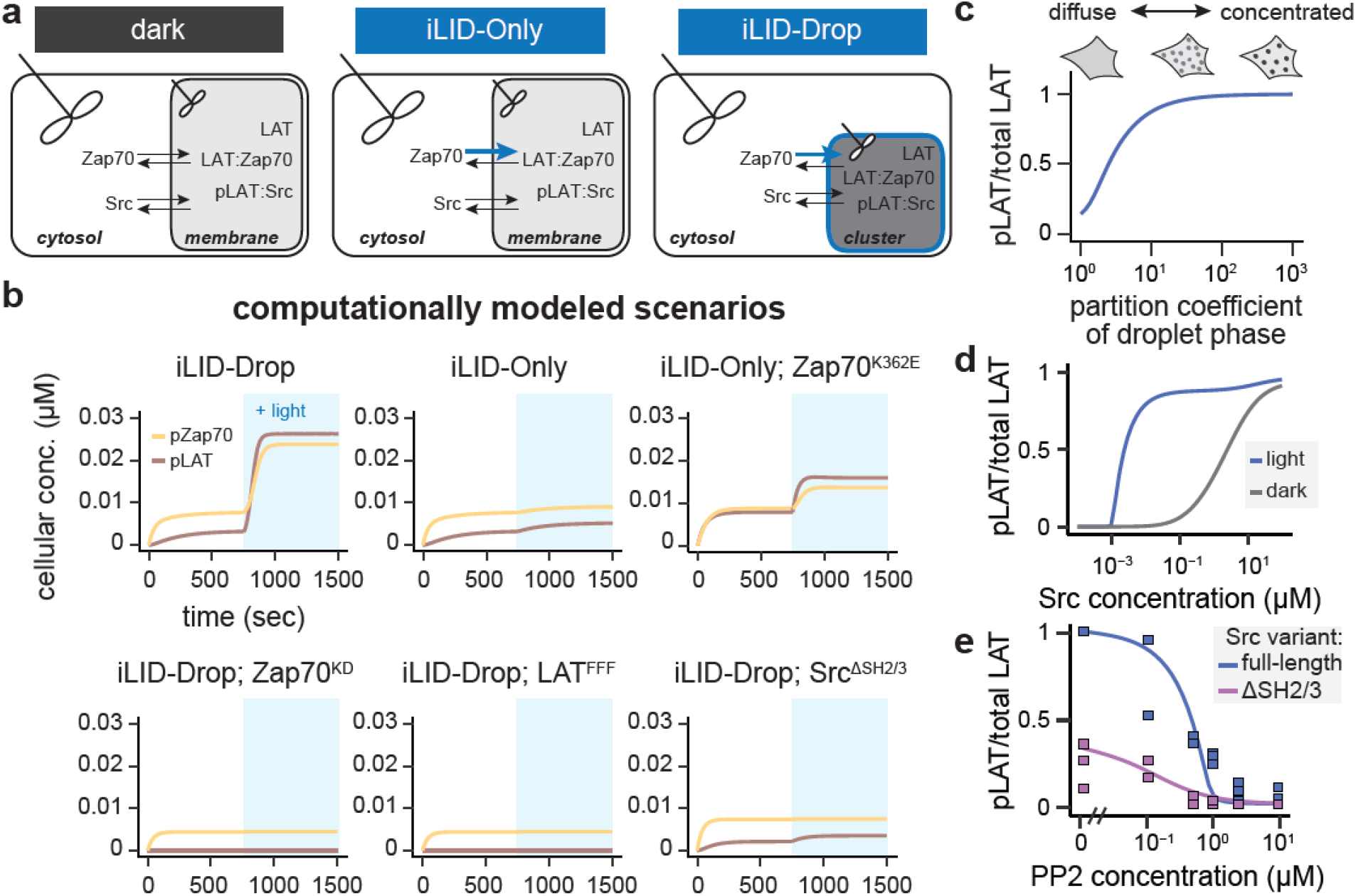
A mathematical model of positive feedback recapitulates sensitivity to clustering state but robustness to SFK concentration. (**a**) Schematic of our mass action kinetic model. In the dark, there are two well-mixed compartments, the cytosol and the membrane, that contain the indicated species. In iLID-Only simulations, stimulation of light leads to Zap70 recruitment to the membrane compartment, while in in iLID-Drop, light stimulation leads to both Zap70 recruitment into the membrane compartment as well as a 10-fold drop in the volume of that compartment. (**b**) Simulated cellular concentrations of pZap70 (yellow line) and pLAT (red line) following light stimulation (blue shading) of indicated optogenetic constructs. (**c**) The modeled ratio of pLAT to total LAT is shown as a function of changes to the “partition coefficient” of LAT *(i.e.,* the extent of the decrease in membrane compartment volume) in light-stimulated iLID-Drop simulations. (**d**) The modeled ratio of pLAT to total LAT is shown as the cellular concentration of Src is varied in iLID-Drop simulations in dark (gray) and light (blue) conditions. (**e**) Model results and experimental data showing the ratio of pLAT to total LAT in response to varying doses of the Src inhibitor PP2 in SYF iLID-Drop fibroblasts expressing either wt Src (blue line) or Src^ΔSH2-3^ (purple line). Straight line shows simulated values while squares show values obtained from Western blots of 3 replicates, except for the 0.1 μM PP2 value which shows 2 replicates. For **e**, modeling results show the iLID-Drop scenario.

We first tested whether this model recapitulated key findings from our experiments. We simulated the model in six experimental scenarios, measuring dark state and light-induced LAT and Zap70 phosphorylation in iLID-Drop, iLID-Only, iLID-Only Zap70^K362E^, iLID-Drop Zap70^KD^, iLID-Drop LAT^FFF^, and iLID-Drop Src^ΔSH2ΔSH3^ cells. In each case, light stimulation was assumed to trigger a 100-fold decrease in the iLID-SspB dissociation constant and a 10-fold increase in LAT-optoDroplet concentration (in the iLID-Drop case only); otherwise parameters were held constant. We observed strong light-induced phosphorylation of LAT and Zap70 in the iLID-Drop but not iLID-Only scenario, with similar kinetics and fold-change in phosphorylation as in our experiments **(Figure 6B**; **Table S2**). The model also matched results from key mutations, showing minimal activity in iLID-Drop simulations harboring kinase-dead Zap70 or non-phosphorylatable LAT. Importantly, our model also requires Src-mediated positive feedback for clusters to trigger signaling. Just as in our experiments, a Src^ΔSH2ΔSH3^ allele that cannot bind phospho-LAT results in an intermediate level of phosphorylation regardless of illumination conditions (**Figure 6B**). The model thus confirms that a clustering-based positive feedback loop is sufficient to quantitatively explain our data across a wide range of experimental conditions.

We next used the model to interrogate the striking combination of sensitivity and robustness revealed by our experiments. It appears that signaling depends sensitively on whether Zap70:LAT complexes are clustered (**Figure 3B-C**), yet the circuit appears to be robust to a ~100-fold variation in Src-family kinase expression (as observed between NIH-3T3 and SYF cells; **Figure S4A**). What degree of LAT clustering is required to trigger a potent signaling response, and over what range of Src concentrations might the circuit function? To address these questions, we first modeled LAT phosphorylation in iLID-Drop cells while varying the degree of light-induced clustering (**Figure 6C**). We observed that signaling increased markedly with the degree of Zap70:LAT clustering, plateauing to a maximum as LAT was concentrated approximately 10-fold above its initial value, well within the range of observed values for protein condensates *in vitro* and in cells^15,49^. In contrast, we observed a strong clustering-induced signaling response even as Src levels were varied across at least two orders of magnitude (**Figure 6D**). Modeling results revealed that both the sensitivity to clustering and robustness to Src concentration absolutely required positive feedback, as simulating the feedback-disconnected Src allele revealed Zap70 and LAT phosphorylation that increased more gradually with Src concentration, and failed to discriminate between clustered and unclustered conditions (**Figure S5A-B**).

As a final probe of the model, we set out to test a prediction in a context not yet measured experimentally: how the signaling module responds to titrating Src activity, not just concentration. To address this question, we simulated a titration of the small-molecule inhibitor PP2 for both wild-type Src and feedback-disconnected Src^ΔSH2ΔSH3^, and then compared to corresponding experimental results. Once again, we found that model and experiment agreed closely, revealing that wild-type Src elicited higher levels of LAT phosphorylation – and signaled effectively across a broader range of PP2 concentrations – than its feedback-disconnected counterpart (**Figure 6E**). Taken together, our computational modeling results confirm that the Zap70/LAT/Src positive feedback circuit can indeed act as a sensitive sensor of protein clustering, while being robust to variation in other cellular parameters (e.g. the concentration or activity state of Src).

## Discussion

Protein phase separation and clustering has been proposed to play a role in a wide variety of cellular functions. But in many cases, it remains possible that phase separation is a consequence, not a cause, of signaling pathway activity. This is particularly when so many signaling proteins engage in weak, multivalent binding interactions that depend on pathway activity (e.g. binding between SH2 domains and pTyr residues). In this study, we set out to determine whether the clustering of two T cell signaling proteins, the kinase Zap70 and its substrate LAT, plays a functional role in modulating downstream signaling. Indeed, we found that Zap70:LAT clustering was sufficient to activate canonical downstream pathways even in non-T cells, whereas a similar number of Zap70:LAT dimers was not. Studies in knockout cell lines and with mutant proteins further revealed the mechanism of cluster-specific signaling: a three-component feedback loop where Src-family kinases bind to phosphorylated LAT, leading to further Zap70 activation (**Figure 7**).

**Figure 7.**
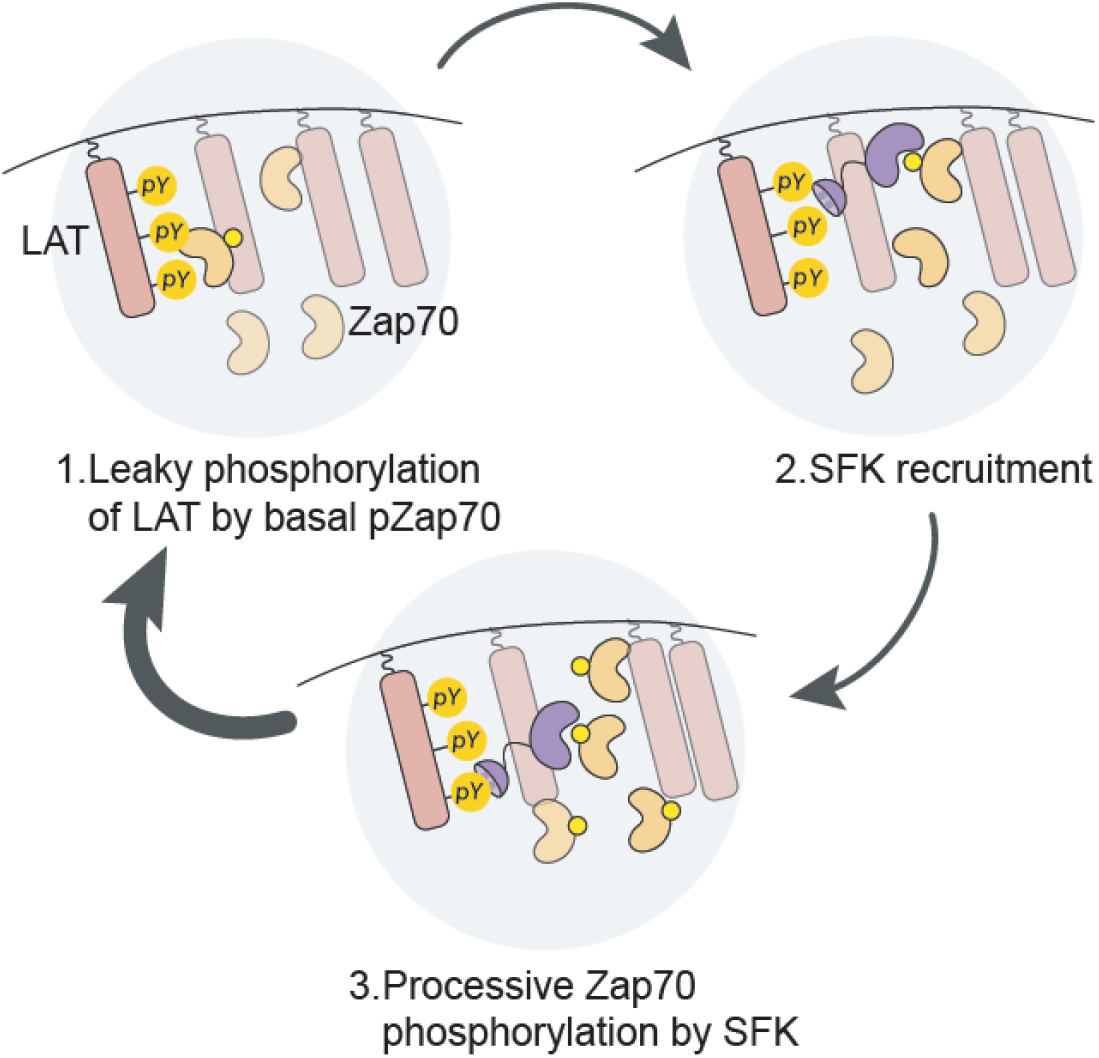
Schematic of the cluster-localized positive feedback loop between pLAT, Src and pZap70. Upon colocalization of LAT and Zap70, an initially low basally-phosphorylated population of Zap70 proteins performs some leaky phosphorylation of LAT. The presence of LAT pTyr residues then enables Src-family kinase (SFK) recruitment and activation through SH2-pTyr and SH3-proline rich motif (PRM) interactions with LAT. Finally, active SFKs phosphorylate additional Zap70 molecules, leading to enhanced Zap70 activity within the cluster and completing the feedback loop.

One major question in cell signaling has been to identify the minimal set of protein components that are required for a particular cellular outcome, such as signaling pathway activation or gene expression. We propose that additional insights can be gained from testing not just which molecular components must be present in the cell, but specifically which must be present in the context of a certain biophysical state (e.g. within a protein condensate or cluster). For example, previous work demonstrated that in Jurkat T cells that other T cell specific proteins such as SLP-76 and GADS are required for downstream signaling^50,51^; yet we observe that fibroblasts expressing neither SLP-76 nor GADS can activate downstream pathways in response Zap70:LAT clustering. It may be that those adaptor proteins are essential for nucleating Zap70:LAT clusters, a function that is provided instead by our optogenetic systems. Separating the creation of a biophysical compartment from signal propagation within it could be of great utility for clarifying the essential functions of components within a signaling pathway.

Cells employ biochemical networks to sense a diverse array of upstream inputs, including extracellular ligands, misfolded proteins, and small molecules. Our study defines a three-component signaling circuit that appears to function as a “condensate detector”. Both experiments and computational modeling reveal that the Zap70/LAT/Src circuit responds strongly to the formation of membrane-localized clusters but not lower-order molecular complexes. Moreover, the system appears to function robustly as other parameters are varied (e.g., the cellular concentration or activity of Src-family kinases). We anticipate that variations of this biochemical circuit may find application in diverse contexts, from biosensors to report on the presence of specific condensates^52^ to synthetic biology studies aiming to engineer novel signaling circuits using designer membraneless organelles^18,19,53^.

Nevertheless, there is still much to do. One clear limitation of the current study is that it leaves open the question about how clustering-based positive feedback affects Zap70 phosphorylation in intact T cells, rather than fibroblasts. Answering this question is complicated by the fact that Zap70:LAT clusters are just one of many distinct types of membrane-associated clusters in the activated T cell: TCR clusters, co-stimulatory clusters and inhibitory clusters^54^. Moreover, Zap70 itself clusters with many other Tyr-rich substrates, including the T cell receptor, and our framework would predict that indeed any of these Tyr-containing substrates could trigger additional feedback phosphorylation of Zap70 via the action of SFK, which may explain why Zap70 phosphorylation remains high in T cells that lack LAT clusters^7,21,35^. The complexity of the native system suggests that much work remains to be done to understand the roles played protein clustering in T cell activation. Reconstitution in fibroblasts presents one possible route to separating these effects, one cluster at a time.

## Acknowledgements

We thank all members of the Toettcher lab for their insights and suggestions throughout the project, particularly Pavithran Ravindran for assistance with figure design. This work was supported by a Vallee Scholars award and NIH grant DP2EB024247 (to J.E.T.), NIH Ruth Kirschstein fellowship F31 AI145218-01 (to E.D.) and NIBB-Princeton postdoctoral fellowship (to E.H.R.). We also thank Dr. Christina Decoste and Katherine Rittenbach of the Princeton Flow Cytometry Core for help with creation of the cell lines used in this manuscript.

## Author Contributions

E.D., and J.E.T. conceived and designed the project and wrote the manuscript. E.H.R. wrote the mathematical model and performed all simulations. E.D. performed all experiments.

## Online Methods

### Plasmids

All plasmids were constructed using inFusion cloning (Clontech) to ligate in a PCR product to a pHR vector that were linearized using backbone PCR.

#### LAT-Zap70 optogenetic constructs

To create iLID-Drop (pHR-TagRFP-SspB-Zap70-P2A-LAT-iLID-FUS^N^-Cry2), we start with the iLID-SspB SOS^cat^ plasmid from Goglia et al. 2020^55^ (Addgene # forthcoming). We removed SOS^cat^ and replaced it with Zap70 from its pDONR plasmid (Addgene # 23387). We removed the CAAX tag and replaced with FUS^N^-Cry2 from myristoylated optoDrop plasmid used in Dine et al. 2018^16^ (Addgene #111507). Finally, we linearized this plasmid via backbone PCR to insert LAT (cDNA obtained from the human ORFeome collection)^56^ between the P2A and iLID sequences in our construct.

We conducted site-directed mutagenesis on Cry2 to make the D387A mutations for the iLID-Only construct. Site-directed mutagenesis was also used to make constitutively active or kinase-dead Zap70 variants as seen in Figures 3 and 4. Site-directed mutagenesis was also to make the point mutants for the experiments displayed in Figure S3. For the LAT^FFF^ iLID-Drop construct, we inserted LAT^FFF^ from Su et al. 2016 (Addgene # 78517)^57^ into the iLID-Drop plasmid. All iLID-Drop and iLID-Only plasmids used in this study have been deposited in Addgene (accession numbers forthcoming).

#### Reporter plasmids

We used pHR-ErkKTR-irFP to monitor activity as we had done previously in Dine et al. 2018^16^ (Addgene # 11510). We used GCaMP6f to monitor calcium activity by performing backbone PCR on a pHR vector and inserting GCaMP6f, obtained by PCR amplification from Addgene plasmid # 10837^30^.

#### SFK plasmids

We performed backbone PCR to linearize the ErkKTR-BFP plasmid from Goglia et al. 2020^55^ (Addgene # forthcoming) and replaced the ErkKTR with a Src-family kinase (SFK) from its respective pDONR plasmid (Src = Addgene # 23934, Fyn = Addgene # 82211 and Lck = Addgene # 82305). Site-directed mutagenesis was then used to create each of the Src variants studied in Figure 5. To make Src^ΔSH2/3^-BFP, we removed the sequence coding for amino acids 83-535 in the original pHR-Src-BFP vector and replaced it with an insert with the sequence coding for amino acids 248-535.

### Cell culture

NIH 3T3, HEK293T (Lenti-X) as well as Src^-/-^ Yes^-/-^ and Fyn^-/-^ (SYF) mouse embryonic fibroblasts (purchased from ATCC) were grown in DMEM supplemented with 10% FBS, 1% L-Glutamine, and penicillin/streptomycin (ThermoFisher Scientific). Cells were maintained on cell culture treated flasks with filter caps (ThermoFisher Scientific) and grown at 37°C with 5% CO_2_.

### Lentivirus production and transduction

Lentivirus was produced as per the protocol we described previously^58^. Briefly, Lenti-X HEK293T cells were plated in a 6-well plate at 20-30% confluency and co-transfected with the appropriate pHR expression plasmid and lentiviral packaging plasmids (pMD2.G and p8.91 – gifts from the Trono lab) using the Fugene HD transfection reagent and manufacturer’s protocols (Promega). Viral supernatants were collected 48-52 hrs after transfection and passed through a 0.45 μm filter.

NIH 3T3, SYF-MEFs and Lenti-X 293T cells to be infected with lentivirus were plated in a 6 well dish at 20% – 40% confluency. 500 μl of filtered virus were added to the cells as was 50 μL of 1 M HEPES. Cells were then grown up and plated in T75 flasks (ThermoFisher Scientific) for cell sorting via FACS Aria as described previously^55^.

### Cell preparation for imaging

For all imaging experiments, cells were plated on black-walled 0.17 mm glass-bottomed 96 well plates (In Vitro Scientific). Prior to cell plating, glass was pretreated with a solution of 10 μg / mL fibronectin in phosphate buffer saline (PBS) for 5 - 60 min (ThermoFisher Scientific). NIH-3T3 and SYF MEFs were allowed to adhere for at least 4 hours in supplemented DMEM. Cells were then switched to starvation media (DMEM + 20 μM HEPES) for 2 hours before imaging. Just prior to imaging 50 μL of mineral oil was added to the top of each well to stop evaporation^59^.

### Time-lapse microscopy

Cells were maintained at 37°C with 5% CO_2_ for the duration of all imaging experiments. Confocal microscopy was performed on a Nikon Eclipse Ti microscope with a Prior linear motorized stage, a Yokogawa CSU-X1 spinning disk, an Agilent laser line module containing 405, 488, 561 and 650 nm lasers, an iXon DU897 EMCCD camera, and 20X air, 40X air, or 60X oil immersion objective lenses.

Due to the fast off-time of our optogenetic constructs, we were only able to image one field of view on our microscope for each experiment, so that the field of view could remain illuminated with blue light in between imaging time points. For every experiment we imaged the ErkKTR with the 650 nm laser, Zap70 localization with the 561 nm laser and GcAMP6f with the 488 nm laser. We acquired these images every 15 sec for 15 min. For the images in Figure 1 and Movies S1-2 we imaged only on the 561 nm laser and did so every 5 sec for 5 min. Between each acquisition for all experiments, we used a 450 nm LED light source (XCite XLED1) delivered through a Polygon400 digital micromirror device (DMD; Mightex Systems) to deliver a constant input of blue light. We set the blue light LED to half its maximal intensity but allowed all the light to pass through the mirrors (no dithering) to provide a strong enough light input for each position imaged.

To collect the data in Figure S4B we imaged the indicated SYF cells with one pulse of 405 nm light to view and measure SFK-BFP expression.

### Cell Lysate Collection

To prepare cells for stimulation and lysis 24 hours prior to experiment cells plated into 6-well dishes at 30% - 40% confluency. The day of experiment cells were checked to be between 60% - 70% confluency. The media was then removed and replaced with 2 ml of starvation media for 2 hrs. Cells were either kept in the dark or stimulated with blue light.

Blue light was delivered via custom-printed boards containing small 450 nm LED bulbs. These boards were placed on top of foil wrapped boxes that were placed in our 37°C incubator. The 6-well dishes containing the cells were then added to the boxes and the blue light board were placed on top of the boards so as too directly stimulate only our cells of interest. Blue light was applied at a constant 5V for 20 min.

Following the 20 min stimulation the media was quickly removed and cells were placed on ice and treated with 120 μl of RPPA lysis buffer (1% Triton X-100, 50 mM HEPES buffer, 150 mM NaCl, 1.5 mM MgCl_2_, 1 mM EGTA, 100 mM NaF, 10 mM sodium pyrophosphate, 1 mM Na3VO4, 10% glycerol, freshly-prepared protease/phosphatase inhibitors). Cell scrapers (Sigma Aldrich) were then used to collect the cells and each lysate was transferred to Eppendorf tubes on ice. Lysates were then spun down at 4°C for 10 min at 13,300 x g. Supernatants were transferred to new tubes where 40 μl 4X NuPAGE LDS Sample Buffer (ThermoFisher Scientific) was added to each, and samples were boiled at 98° C for 5 min.

### Western Blotting

Samples were then run on a gel for western blotting done as described previously in Goglia et al. 2020^55^. Primary antibodies used in this study are as follows: rabbit-anti-pY319-Zap70 (Cat # 2717S), mouse-anti-Zap70 (Cat # 2709S), rabbit-anti-pY191-LAT (Cat # 3584s), rabbit-anti-GAPDH (Cat# 2118s), rabbit-anti-ppErk (Cat # 4370s), mouse-anti-Erk (Cat # 4696s), mouse-anti-Src (Cat # 2210s) all purchased from Cell Signaling Technologies, and rabbit-anti-pY132-LAT (Cat # 44-244) and mouse-anti-LAT (Cat # 14-9967-82) purchased from Invitrogen. All primary antibodies were used at 1:1,000 dilution, except for anti-GAPDH, which was used at 1:2,500. Fluorescent secondary antibodies, 800CW goat anti-rabbit (Cat # 926-32211) and IRDye 680RD goat anti-mouse (Cat # 926-68070) were purchased from Li-Cor and used at a 1:10,000 dilution.

Blots were then imaged on imaged on a Li-Cor Odyssey CLx imaging system, and images were analyzed using imageJ software (FIJI) to calculate pixel intensities for all bands of interest. Pixel intensities from phospho-species antibodies were divided by the corresponding total species value as indicated in each figure. For the blot measuring LAT or Src expression levels in Figures S1B and S4A respectively, the band intensities in the 680 channel (anti-LAT or anti-Src) was divided by the corresponding values in the 800 channel (anti-GAPDH). Plotting and statistical analysis for all blots was performed using GraphPad Prism 8. Unpaired student’s T tests were used to compare dark and light conditions for each different cell line or drug treatment.

### Drug Additions

To inhibit SFK activity PP1 and PP2 (Millipore Sigma) were reconstituted at a concentration 10 mM in DMSO and kept at −20°C. Immediately prior to cell stimulation with blue light (or darkness) PP1 and PP2 were diluted a total of 1,000 fold in starvation media (for a final concentration of 10 μM) and added to cells to acutely inhibit SFK activity.

### Image analysis

All image analysis was performed in ImageJ (FIJI) and all plots were generated in Graphpad Prism 8.

#### Measuring Zap70 Cytosolic and Membrane Intensity

To track Zap70 cytosolic intensity, we imaged through the central z plane of cells to track TagRFP intensity in the cytosol. We then hand selected regions of cytosol from each cell and measured the mean TagRFP fluorescence values in that region for every time point. We background subtracted every measured time point and normalized the intensity values of each cell to the initial (dark state) TagRFP cytosolic intensity to generate the plot shown in Figure 1E. Measuring the mean TagRFP intensity of hand-drawn cytosolic regions of 25 iLID-Drop and iLID-Only cells in the dark state was used to generate the box and whiskers plot shown in Figure S1C.

To track the coefficient of variation of Zap70 membrane intensity, we imaged cells through their membrane plane, allowing us to track TagRFP intensity at the membrane. We measured both mean TagRFP intensity and the standard deviation of the intensity in hand-drawn regions of interest on each cell membrane for each time point. We then divided the standard deviation by the mean intensity at each time point for every cell. We normalized these CV values to the initial (dark state) CV value to generate the plot shown in Figure 1F. Measuring the mean BFP intensity of hand-drawn regions of the membrane of 20 cells of each of the indicated SYF cell lines was used to generate the box and whiskers plot shown in Figure S4B.

#### Measuring KTR and GCaMP values

For KTR analysis equivalent nuclear or cytoplasmic regions were tracked over time by hand annotation. We then measured the mean fluorescent intensity in each annotated region for every time point. We then background subtracted every measured time point and plotted the cytoplasmic/nuclear ratio for each time point. The Area Under the Curve (AUC) was calculated by subtracting the initial (dark state) cytosolic to nuclear ratio from the value at each time point and summing up all those differences for each of the 61 time points.Non-paired student’s T tests were used to compare dark and light conditions for each different cell line or drug treatment.

For GCaMP analysis, a small area was drawn in a randomly chosen cytoplasmic region of each cell. Mean fluorescent intensities were measured and background subtracted as above. Values were then normalized to the minimum value found in each cell’s individual trace. Graphs were generated as was done for ErkKTR. AUC and statistical testing was done as for the ErkKTR calculations.

### Computational Model

Our computational model consists of three species: LAT, Src, and Zap70 that appear in two cellular compartments: the membrane/cluster compartment and the cytosol. LAT resides in the membrane compartment, and Src and Zap70 reside in the cytosol and can diffuse freely into the membrane compartment. We used mass action kinetics to describe phosphorylation of Zap70 and LAT in the dark state and after illumination with blue light under several scenarios (**Table S2**). We also compared the steady state extent of phosphorylation given by our model to the experimentally measured values (**Table S1**) and parameters given in **Table S3**.

To simulate the formation of LAT clusters upon illumination, we decreased the volume of the membrane compartment by a factor, **K**, such that:

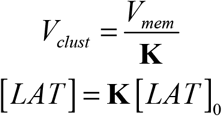

Additionally, Zap70 binds to LAT by an iLID/SspB interaction upon illumination. We assumed that diffusion and binding is much faster than phosphorylation and can therefore be approximated to be at equilibrium and that the cytosolic concentration of Zap70 remains constant. Furthermore, we assumed that Zap70 will also freely diffuse into the clusters, therefore:

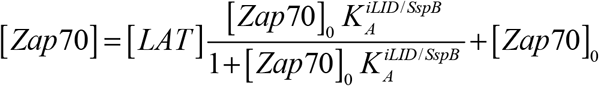

Src can bind to phosphorylated LAT through a SH2/pY interaction and this binding releases autoinhibition. As before, we approximated binding to be at equilibrium and assumed that the cytosolic concentration of Src remains constant. Src will also freely diffuse into the cluster, remaining in an inhibited state, therefore:

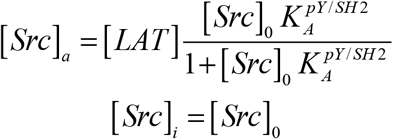

We modeled Zap70 phosphorylation by Src using Michealis-Menten kinetics. In the cluster, Zap70 will be rapidly phosphorylated by active Src and slowly phosphorylated by autoinhibited, inactive Src. In the cytosol, Zap70 will be phosphorylated by inactive Src. Furthermore, we assumed that Zap70 undergoes constitutive dephosphorylation following first order kinetics:

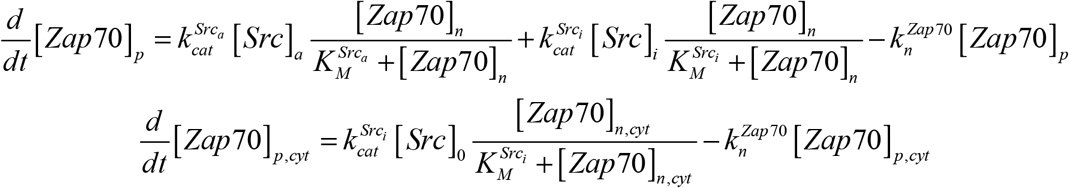

Finally, we modeled phosphorylation of LAT by pZap70. For simplicity, we considered only one phosphorylatable tyrosine on LAT. We allowed LAT to be phosphorylated by two distinct mechanisms: (1) phosphorylation by pZap in the cluster following Michaelis-Menten kinetics, and (2) preferential phosphorylation of LAT by pZap70 that is bound to it by an iLID—SspB interaction following first order kinetics. Additionally, we assumed that LAT undergoes constitutive dephosphorylation following first order kinetics.

LAT that is not bound to Zap70 can only be phosphorylated by the first mechanism, therefore:

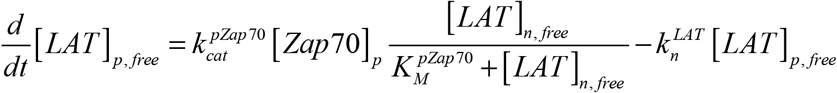

LAT that is bound to Zap70 may be phosphorylated by either phosphorylation mechanism, therefore:

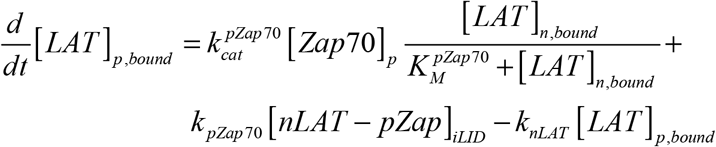

In the preceding equation, [*nLAT-pZap*70]_*iLID*_ is the pool of iLID:SspB bound LAT:Zap70 complexes that consist of nonphosphorylated LAT and phosphorylated Zap70, given by:

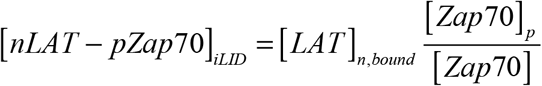

PP2 was modeled as a non-competitive inhibitor, therefore, the catalytic rate constants for active and inactive Src were scaled by:

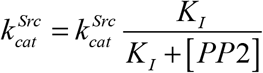

All simulations were performed in MATLAB version R2020a, using ode23 to solve the differential equations. Graphs generated from the model were plotted in R Studio version 1.1.456.

