## Supplementary Figures and Tables for "Clustering-based positive feedback between a kinase and its substrate enables effective T-cell receptor signaling"

**This PDF file includes:**

| | |
|---|---|
| Supplementary Figures 1-5 | Pages 2-7 |
| Supplementary Tables 1-3 | Pages 8-10 |
| Supplementary Movie 1-5 Legends | Page 11 |
| Supplementary References | Page 12 |

### Supplementary Figures and Legends

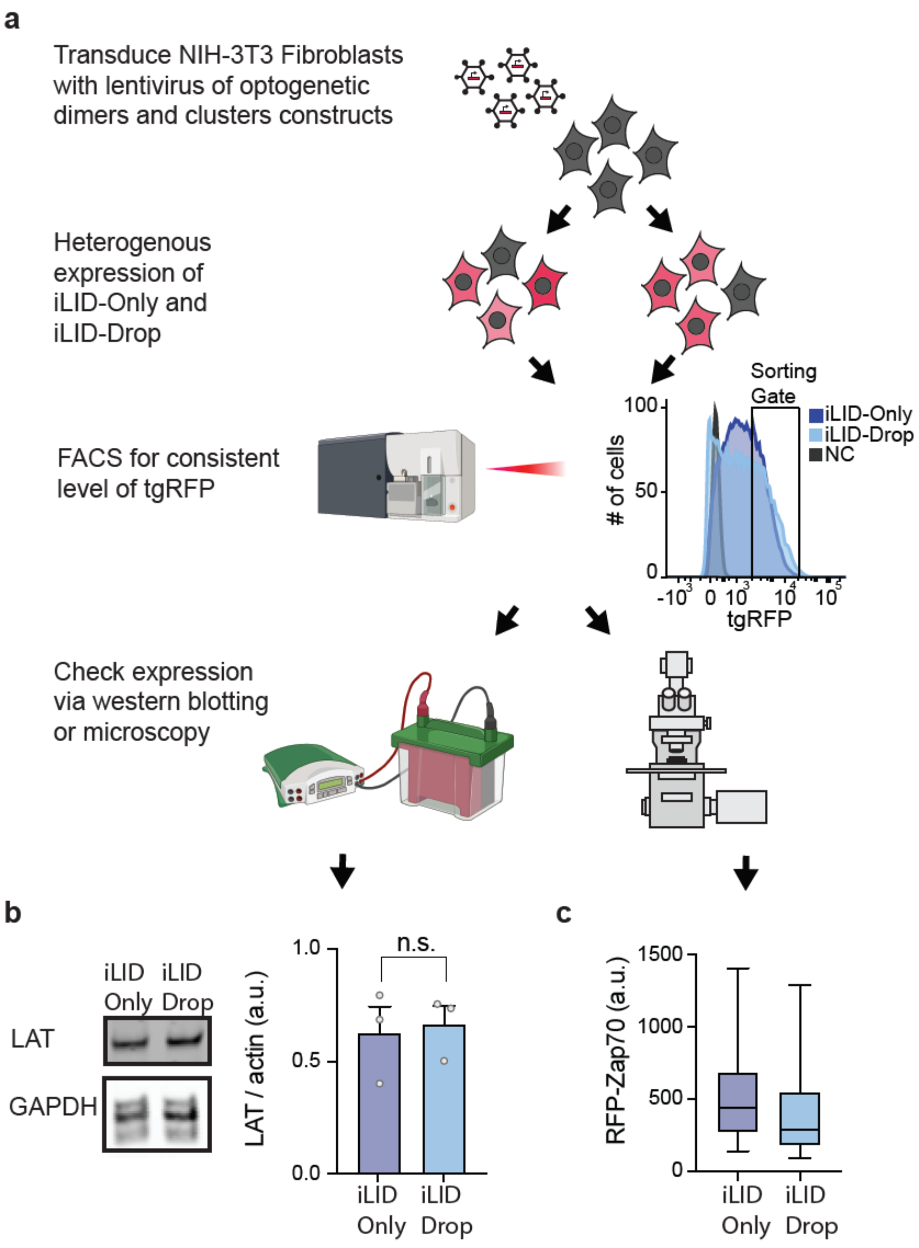

**Figure S1: Ensuring that iLID-Only and iLID-Drop optogenetic constructs are expressed at similar levels in NIH-3T3 cell lines.** (**a**) Flow chart outlining process of creating the iLID-Only and iLID-Drop expressing cell lines used throughout this paper. Plain NIH-3T3 fibroblasts were infected with lentivirus containing constructs depicted in Figure 1B. These stably infected cells were then sorted using a BD systems FACS Aria. The same gate for TagRFP expression were used for both the iLID-Only and iLID-Drop expressing cell lines, as shown. These stable, sorted cell lines were then used for the microscopy and western blotting experiments done in later experiments and to confirm similar levels of expression for both constructs. (**b**) Western blot and quantification of LAT normalized to a loading control (GAPDH) in iLID-Only and iLID-Drop expressing NIH-3T3 cell lines. Graphs display mean, SEM (error bars) and independent biological replicate (dots). $p > 0.05$ from an unpaired Student's T test. (**c**) Box and Whisker plots showing TagRFP (Zap70) fluorescence for iLID-Only and iLID-Drop cells. Boxes represent $25^{th} - 75^{th}$ percentile with line in the middle representing the mean and whiskers show minimum and maximum. $n = 25$ cells for both conditions.

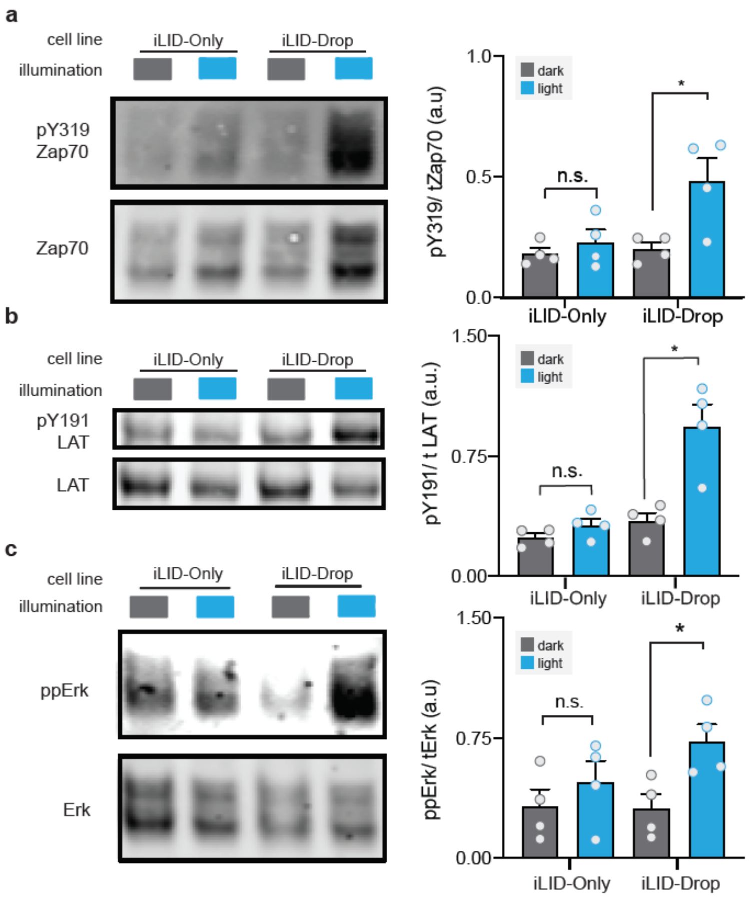


**Fig S2: Induction of optogenetic Zap70:LAT clusters leads to successful signaling in HEK-293T cells.** (**a**) Western blot and quantification for pY319-Zap70 in HEK293T expressing iLID-Only and iLID-Drop cells. For all experiments shown in this figure, cells were either kept in dark (gray bar) or stimulated with blue light for 20 minutes (blue bar). For quantification, dark and light were each compared using the student's t test for all experiments. Graphs display mean, SEM (error bars) and independent biological replicate (dots). * = $p < 0.05$ (**b**) Western blot and quantification for pLAT Y191 in HEK293T expressing iLID-Only and iLID-Drop cells. (**c**) Western blot and quantification of ppERK in HEK293T expressing iLID-Only and iLID-Drop cells.

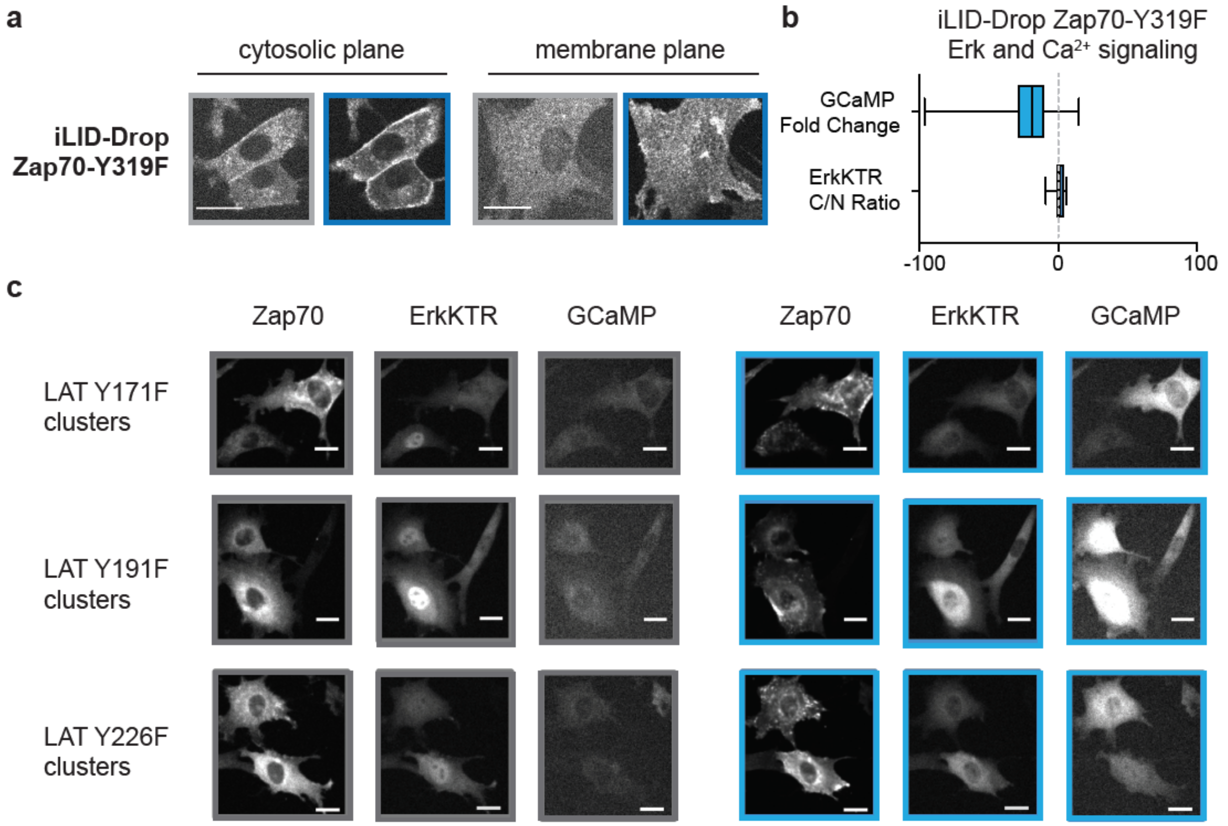


**Figure S3: Zap70 phosphorylation is driven by phosphorylation of all three LAT tyrosines.** (**a**) Images of TagRFP localization in NIH-3T3 cells expressing Zap70-Y319F iLID-Drop. Images of Tag-RFP Zap70-Y319F were taken at two different planes (cytosolic and membrane) with spinning disk confocal imaging. Gray border indicates images taken prior to blue light illumination, and blue border indicates images taken following 5 minutes of stimulates. Note images underwent contrast enhancement in ImageJ to account for the increase in brightness of TagRFP that occurs when illuminated with blue light. Scale bars = 20 µm. (**b**) Box and Whisker plots showing Area Under the Curve (AUC) for cytoplasmic/nuclear ratios of ErkKTR-irFP and GCaMP fluorescence in Zap70-Y319F iLID-Drop expressing NIH-3T3 cells. Boxes represent $25^{th}$ – $75^{th}$ percentile with line in the middle representing the mean and whiskers show minimum and maximum. n ≥ 20 cells from 2 different experiments. (**c**) Representative images showing localization of Zap70, ErkKTR translocation and GCaMP fluorescence for NIH-3T3 cell lines expressing iLID-Drop with single LAT Y → F mutations. Gray squares around images show cells pre-light stimulation and blue squares around show cells during blue light stimulation (10 minutes for Zap70 and ErkKTR and 3 minutes for GCaMP). Note Zap70 images were autoscaled in imageJ to account for the increase in brightness of TagRFP that occurs when illuminated with blue light. Scale bar = 10 µm.

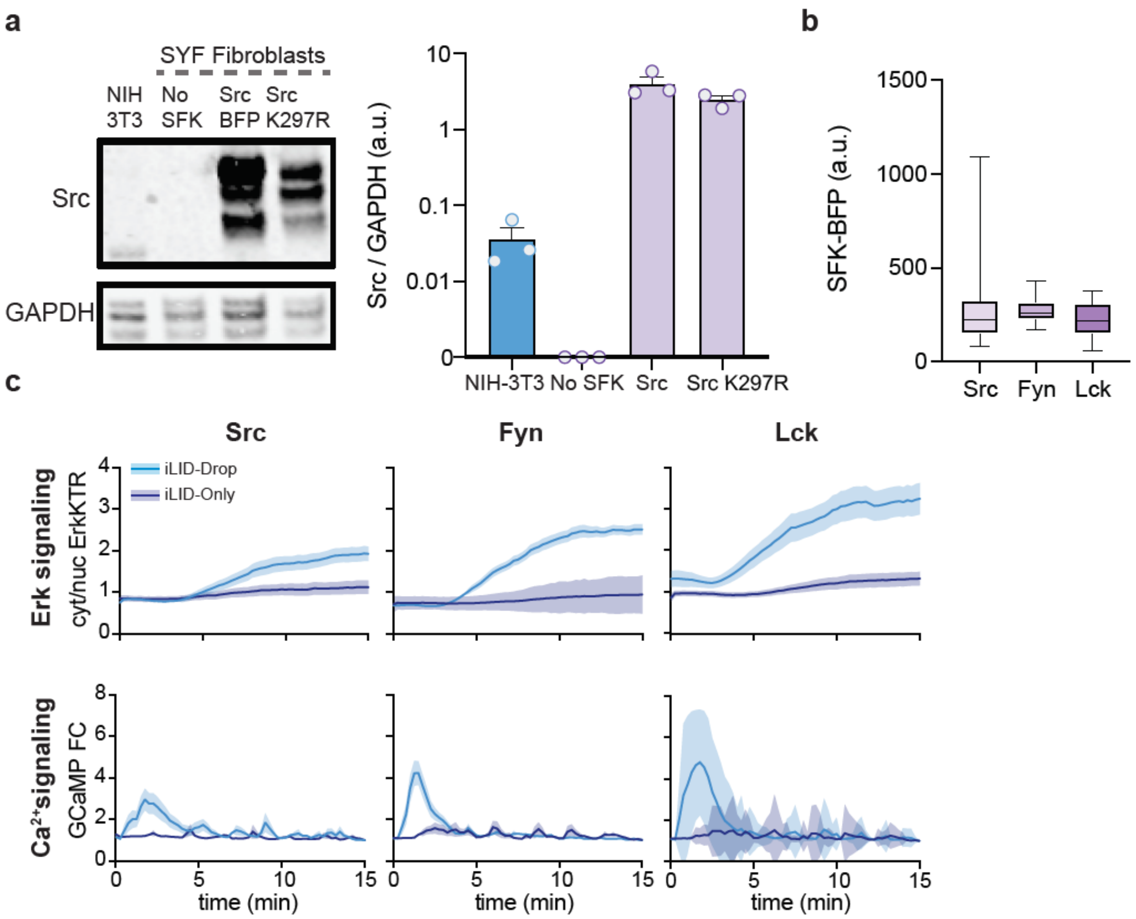


**Figure S4: SFK expression levels in engineered SYF-MEF cell lines.** (**a**) Western blot and quantification of Src normalized to a loading control (GAPDH) in NIH-3T3, SYF, SYF + SRC-BFP and SYF + Src$^{K297R}$-BFP cell lines. Graphs display mean, SEM (error bars) and independent biological replicate (dots). (**b**) Quantification of fluorescence in 405 nm channel (SFK-BFP) post-FACS sorting for SYF-MEF cell lines different SFKs. n ≥ 20 cells for each cell line. (**c**) Traces of ErkKTR cytoplasmic/nuclear ratio or GCaMP fluorescence for different SYF-MEF cell lines expressing iLID-Only and iLID-Drop constructs. Traces show mean with shaded area representing the SEM. n ≥ 20 cells for every condition.

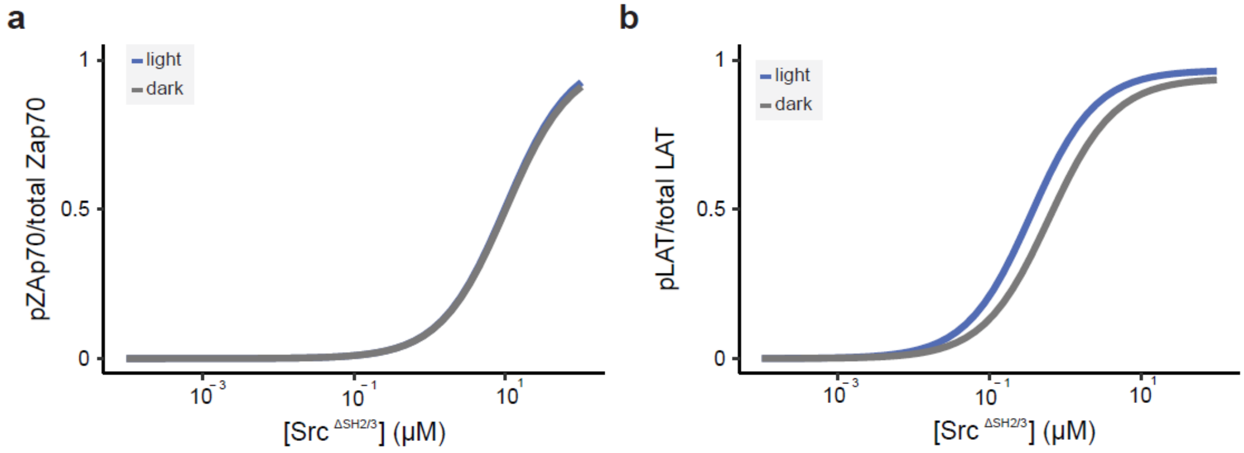


**Figure S5: Zap70 and LAT phosphorylation levels display little difference between dark and light stated in simulated cells expressing iLID Drop and Src$^{\Delta SH2\text{-}3}$**. (**a**) Ratio of pZap70 to total Zap70 in computer simulations with varied Src$^{\Delta SH2\text{-}3}$ concentrations for iLID-Drop cells in dark and light states. (**b**) Ratio of pLAT to total LAT in computer simulations with varied Src$^{\Delta SH2\text{-}3}$ concentrations for iLID-Drop cells in dark and light states.

| Output | Model | Experiment |
|---|---|---|
| $pLAT_{lit}/pLAT_{dark}$ iLID-Only | 1.68 | 1.47 |
| $pZap70_{lit}/pZap70_{dark}$ iLID-Only | 1.19 | 1.02 |
| $pLAT_{lit}/pLAT_{dar}$ iLID-Drop | 10.52 | 6.82 |
| $pZap70_{lit}/pZap70_{dark}$ iLID-Drop | 3.81 | 3.29 |
| $pLAT_{lit}/pLAT_{dark}$ iLID-Only Zap70-K362E | 2.02 | 3.66 |
| $pZap70_{lit}/pZap70_{dark}$ iLID-Drop LAT FFF | 1.02 | 1.3 |
| $pZap70_{lit}/pZap70_{dark}$ iLID-Drop Zap70 KD | 1.02 | 1.3 |
| $pZap70_{lit}/pZap70_{dark}$ iLID-Drop Src ΔSH2 ΔSH3 | 1.02 | 1.38 |

**Table S1: Comparison of model outputs to experimentally measured values. Experimental pLAT values are for Y191.**

| Scenario | Description |
|---|---|
| iLID-Drop | The base scenario. |
| iLID-Only | Blue light insensitive Cry2 on LAT modeled by setting **K** to 1 in the lit state. |
| Zap70-K362E | Weakly constitutively active Zap70 modeled by allowing both phosphoylated and non-phosphorylated Zap70 to phosphorylate LAT with lower rate constant and higher Michelais constant than WT pZap70. |
| Zap70 KD | Kinase dead Zap70 modeled by setting the rate constant for phosphorylation of LAT by pZap70 to 0. |
| LAT FFF | LAT Y to F mutation modeled by setting the pLAT/Src association constant to 0 ($K_D^{pY/SH2}$ equal to Inf.) and the catalytic rate for phosphorylation of LAT by pZap70 to 0. |
| ΔSH2 ΔSH3 | Src ΔSH2 ΔSH3 mutation modeled by setting the pLAT/Src association constant to 0 ($K_D^{pY/SH2}$ equal to Inf.), increasing the rate constant for phosphorylation of Zap70 by non-bound Src, and reducing the cellular concentration of Src. |

**Table S2: Description of computationally modeled scenarios.**

| Parameter | Description | Value |
|---|---|---|
| $[LAT]_0$ | Membrane-localized compartment concentration of LAT | 0.1 µM |
| $[Zap70]_0$ | cytosolic concentration of Zap70 | 1.5 µM |
| $[Src]_0$ | cytosolic concentration of Src | 0.15 µM<br>0.05 µM (ΔSH2 ΔSH3) |
| **K** | LAT concentration fold change upon clustering | 1 (dark)<br>10 (lit, ilID-Drop)<br>1 (lit, iLID-Only) |
| $K_D^{iLID/SspB}$ | iLID/SspB dissociation constant | 13 µM (dark)<br>0.13 µM (lit, iLID-Drop)[1]<br>0.13 µM (lit, iLID-Only) |
| $K_D^{pY/SH2}$ | pLAT/Src dissociation constant | 0.01 µM[2,3]<br>Inf. µM (ΔSH2 ΔSH3)<br>Inf. µM (LAT FFF) |
| $k_{cat}^{Src\text{-}a}$ | Catalytic rate constant for phosphorylation of Zap70 by active Src | 1 $s^{-1}$ |
| $K_M^{Src\text{-}a}$ | Michaelis constant for phosphorylation of Zap70 by active Src | 100 µM |
| $k_{cat}^{Src\text{-}i}$ | Catalytic rate constant for phosphorylation of Zap70 by autoinhibited Src | 0.05 $s^{-1}$<br>0.25 $s^{-1}$ (ΔSH2 ΔSH3) |
| $K_M^{Src\text{-}i}$ | Michaelis constant for phosphorylation of Zap70 by autoinhibited Src | 100 µM |
| $k_{cat}^{pZap70}$ | Catalytic rate constant for phosphorylation of LAT by pZap70 | 1 $s^{-1}$<br>0.25 $s^{-1}$ (Zap70 K362E)<br>0 $s^{-1}$ (LAT FFF)<br>0 $s^{-1}$ (Zap70 KD) |
| $K_M^{pZap70}$ | Michaelis constant for phosphorylation of LAT by pZap70 | 10 µM<br>200 µM (Zap70 K362E) |
| $k^{pZap70}$ | Rate constant for preferential phosphorylation of LAT by tethered pZap70 | 0.05 $s^{-1}$<br>0.0125 $s^{-1}$ (Zap70 K362E)<br>0 $s^{-1}$ (LAT FFF)<br>0 $s^{-1}$ (Zap70 KD) |
| $k_n^{LAT}$ | Rate constant for constitutive dephosphorylation of LAT | 0.01 $s^{-1}$ |

| | | |
|---|---|---|
| $k_n^{Zap70}$ | Rate constant for constitutive dephosphorylation of Zap70 | 0.025 $s^{-1}$ |
| $V_{cell}$ | Volume of the cell | 8000 $\mu m^3$ |
| $V_{mem}$ | Volume of the membrane-localized compartment | 2400 $\mu m^3$ |
| $K_I^{PP2}$ | Src/PP2 dissociation constant (equal to IC50 when modeled as a non-competitive inhibitor) | 0.1 μM |

**Table S3: Description and values of model parameters.**

### Supplementary Video Legends

**Movie S1.** Confocal imaging of NIH-3T3 cells expressing iLID-Only (left-hand side) or iLID-Drop constructs (right-hand side). Images were acquired with RFP imaging settings (561 nm excitation) every 5 sec for 5 min to visualize TagRFP-SspB-Zap70 localization. Images were taken in the Z slice through the middle of the cell to observe cytosolic depletion of Tag TagRFP-SspB-Zap70 upon blue light stimulation (marked with blue line). Note that first image was copied 10 times to give a clear view of the initial dark state and that all images underwent process enhancement to account for rise in TagRFP fluorescence that occurs when stimulated with blue light. Scale bar = 20 μM. Related to Figure 1.

**Movie S2.** Confocal imaging of NIH-3T3 cells expressing iLID-Only (left-hand side) or iLID-Drop constructs (right-hand side). Images were acquired with RFP imaging settings (561 nm excitation) every 5 sec for 5 min to visualize TagRFP-SspB-Zap70 localization. Images were taken in the Z slice at the plasma membrane plane of the cell to observe formation of membrane clusters of Zap70 and LAT in iLID-Drop cells upon blue light stimulation (marked with blue line). Note that first image was copied 10 times to give clear view of the initial dark state and that all images underwent process enhancement to account for rise in TagRFP fluorescence that occurs when stimulated with blue light. Scale bar = 20 μM. Related to Figure 1.

**Movie S3.** Confocal imaging of NIH-3T3 cells expressing iLID-Drop optogenetic construct along with ErkKTR-irFP and GCaMP6f. Images from left to right show changes in localization and clustering of TagRFP-SspB-Zap70, nuclear-cytosolic shuttling of ErkKTR-irFP and oscillations in GCaMP fluorescence upon blue light stimulation (marked with blue line). Images were taken using RFP (561 nm) irFP (640 nm) and GFP (488 nm) settings. Note that the first frame of the movie was copied 10 times to give a clear view of the initial dark state and that the RFP channel underwent process enhancement in imageJ to account to account for rise in TagRFP fluorescence that occurs when stimulated with blue light. Scale bar = 20 μM. Related to Figure 2.

**Movie S4.** Confocal imaging of NIH-3T3 cells expressing iLID-Only optogenetic construct along with ErkKTR-irFP and GCaMP6f. Images from left to right show changes in localization of TagRFP-SspB-Zap70, no change in ErkKTR-irFP localization and dim GCaMP fluorescence even upon blue light stimulation (marked with blue line). Images were taken using RFP (561 nm) irFP (640 nm) and GFP (488 nm) settings. Note that the first frame of the movie was copied 10 times to give a clear view of the initial dark state and that the RFP channel underwent process enhancement in imageJ to account to account for rise in TagRFP fluorescence that occurs when stimulated with blue light. Scale bar = 20 μM. Related to Figure 2.

**Movie S5.** Confocal imaging of NIH-3T3 cells expressing iLID-Only Zap70 K362E optogenetic construct along with ErkKTR-irFP and GCaMP6f. Images from left to right show changes in localization of TagRFP-SspB-Zap70, nuclear-cytosolic shuttling of ErkKTR-irFP and oscillations in GCaMP fluorescence upon blue light stimulation (marked with blue line). Images were taken using RFP (561 nm) irFP (640 nm) and GFP (488 nm) settings. Note that the first frame of the movie was copied 10 times to give a clear view of the initial dark state and that the RFP channel underwent process enhancement in imageJ to account to account for rise in TagRFP fluorescence that occurs when stimulated with blue light. Scale bar = 20 μM. Related to Figure 3.
